# Miniscope Processing Suite (MPS): An Intuitive, No-Code, Scalable Pipeline for Long-Duration Calcium Imaging

**DOI:** 10.64898/2025.12.01.691517

**Authors:** Ari Peden-Asarch, Meredith Weinstock, Kevin R. Coffey, John F. Neumaier

## Abstract

Miniscope calcium imaging provides a unique window into the activity of neurons during behavior while enabling spatial localization of individual cells across time. Despite its potential to revolutionize *in vivo* imaging alongside the rise of optogenetic tools, miniscopes remain underutilized. This gap may stem from the lack of an easy-to-use preprocessing software tailored to miniscope data. To address this need, we developed the first no-code end-to-end scalable pipeline for preprocessing large miniscope recordings: the Miniscope Processing Suite (MPS). MPS is the first implementation of Constrained Non-negative Matrix Factorization (CNMF) with Nonnegative Double Singular Value Decomposition (NNDSVD) initialization, multi-lasso segmentation, and parallelized temporal and spatial updates, enabling analysis of recordings exceeding three hours on a standard hardware. We tested a large dataset consisting of 28 operant behavior sessions (77 hrs, 7.26 TB of video data) on a single workstation. MPS completed the analysis in 55.6 hrs, (averaging 0.72 minute of processing time per minute of recording), which is 10-20X faster than traditional pipelines. Packaged as an easy downloadable plug-and-play software with a graphical user interface and requiring no coding or Git experience, MPS lowers the barrier to miniscope analysis while improving on current methodologies.

## Introduction

Miniaturized one-photon fluorescence microscopes (“miniscopes”) have revolutionized *in vivo* neural imaging by enabling single-neuron resolution recordings in freely behaving animals (***Ghosh et al.*** (***2011***); ***Ziv et al.*** (***2013***)). Unlike tethered two-photon setups, miniscopes are lightweight enough to be head-mounted on rodents, permitting neural activity to be monitored during natural behaviors without restraining the subject (***Aharoni and Hoogland*** (***2019***)). Moreover, through implanted microendoscopic gradient-index (GRIN) lenses, researchers can perform chronic imaging of previously inaccessible cell populations deep in the brain over weeks to months (***Ziv and Ghosh*** (***2015***); ***Zhou et al.*** (***2018***)). As a result, researchers can theoretically track the same neurons across multiple sessions and then study evolving neuronal dynamics (***Ziv et al.*** (***2013***); ***Cai et al.*** (***2016***)). Additionally, recent hardware advances have further enhanced miniscope capabilities with newer designs that support simultaneous dual-wavelength imaging (e.g. recording green and red indicators in parallel) to distinguish cell populations or combine activity reporters with stable reference markers (***Barbera et al.*** (***2024***); ***Dong et al.*** (***2025***)). Thus, miniscopes integrate single-cell resolution with behavioral compatibility and chronic repeatability and can be implemented in conventional behavioral procedures over repeated days, making them a powerful tool for neuroscience.

The proliferation of miniscope users created a pressing need for user-friendly, scalable analysis pipelines. One-photon calcium imaging data pose special challenges: signals are often confounded by movement artifacts, background neuropil autofluorescence, and overlapping neuronal footprints, making automated source extraction difficult (***Zhou et al.*** (***2018***)). Several computational pipelines have emerged to address these issues, each offering valuable algorithms but with notable practical limitations. To our knowledge, the longest continuous miniscope recordings processed to date span ∼45–60 minutes per session across longitudinal studies extending up to several months (***Ziv et al.*** (***2013***); ***Cai et al.*** (***2016***); ***Sheintuch et al.*** (***2017***); ***Gonzalez et al.*** (***2019***); ***Jennings et al.*** (***2019***); ***Chen et al.*** (***2023***); ***Zhou et al.*** (***2020***); ***Guo et al.*** (***2023***); ***Wirtshafter and Disterhoft*** (***2022***); ***Blair et al.*** (***2023***); ***Vergara et al.*** (***2025***)). Theoretically, current pipelines can handle longer behavioral experiments by chunking datasets into separate files, but this can compound errors in motion correction and fails to preserve the true longitudinal stability of neuronal identities over time - a limitation stemming largely from the constraints of existing software. CaImAn is a widely used open-source library that implements motion correction and constrained non-negative matrix factorization (CNMF) for source separation and achieves near-human accuracy in detecting active neurons (***Giovannucci et al.*** (***2019***)). However, CaImAn and related packages like CNMF-E (an optimized CNMF for microendoscope data) require users to write code or scripts in MATLAB/Python and adjust parameters programmatically and demand navigating a difficult setup process across Integrated Development Environments (IDEs), virtual environments, and multiple package managers (***Giovannucci et al.*** (***2019***); ***Zhou et al.*** (***2018***)). Other pipelines such as EXTRACT (a GPU-accelerated cell extraction algorithm primarily designed for two-photon datasets, lacking the robust motion correction needed for miniscope recordings) and MIN1PIPE (which introduced morphological background removal and seeded CNMF initialization) likewise demand considerable coding or technical expertise to operate (***Lu et al.*** (***2018***); ***Inan et al.*** (***2021***)). More recently, Minian/CaliAli provided an “open-source miniscope analysis pipeline” with interactive visualization of parameters to guide users with minimal computational experience (***Dong et al.*** (***2022***); ***Vergara et al.*** (***2025***)). These softwares reduce memory usage by leveraging Dask’s out-of-core computation, allowing analysis to run on consumer-grade hardware. Nonetheless, this strategy comes with trade-offs: memory is fragmented across processes and not released until the script or IDE is fully closed, and long recordings eventually collapse into a single oversized task graph. For example, a typical 600 × 600 field of view recorded for 95,000 frames (∼ 3 hours) already produces 34.2 × 10^9^ data points. When multiplied across ∼ 1,000 components for matrix factorization, this balloons to 34.2 × 10^12^ elements – requiring more than ∼ 270TB of memory if handled naively in 64-bit precision. In practice, this Dask graph attempts to juggle this scale of computation, but since the entire dataset is put into memory at once at a key CNMF related step, the processing of datasets of this size is impossible. Users must also still launch Jupyter notebooks or scripts, rather than relying on a stable standalone GUI. (***Dong et al.*** (***2022***)) Additionally, even given a machine with ∼ 512GB of RAM, the largest video size successfully run before saturating memory and causing a crash was a 65 minute video with 600 × 600 pixels with MIN1PIPE. In practice, running and maintaining pipelines of similar scale often demands deep computer science expertise (managing virtual environments, CUDA dependencies, and package conflicts) that many biology labs rightfully lack. Lastly, deep-learning–based ROI detection has emerged in recent software; MPS is complementary to these approaches and supports integration of lab-specific models (***Tian et al.*** (***2025***)). Therefore, MPS directly addresses this gap by integrating state-of-the-art algorithms into a stable, point-and-click platform engineered to process multi-hour recordings on everyday workstations. No existing solution fully combines advanced calcium imaging algorithms with an accessible, graphical user interface (GUI) that can reliably handle large datasets without overwhelming standard laboratory hardware. As many labs find data processing to be a bottleneck and lack the coding expertise to take full advantage of miniscope technology, it is not surprising then to see that miniscopes have been underutilized by the field at large.

In this paper, we introduce the Miniscope Processing Suite (MPS), a GUI-based, end-to-end calcium imaging pipeline optimized for long-duration miniscope recordings. MPS builds on the gold-standard CNMF framework (***Pnevmatikakis et al.*** (***2016***)), but adapts it for long-duration miniscope data. Beyond Non-negative Double Singular Value Decomposition (NNDSVD) initialization and custom multi-lasso regularization, MPS introduces (i) robust preprocessing with neuropil subtraction, denoising, and cross-correlation motion correction; (ii) interactive regions of interest (ROI) cropping to reduce data load by ∼70%; (iii) watershed-based segmentation with merging/validation to avoid duplicate or fragmented ROIs; (iv) parallelized temporal and spatial CNMF updates with out-of-core Dask execution, enabling stable analysis of multi-hour datasets; (v) automated artifact detection (e.g., erroneous frames and line-splitting events); and (vi) a built-in GUI Data Explorer for final quality control. These combined improvements allow MPS to scale where prior pipelines fail, while remaining accessible to non-programmers. Additionally, The pipeline is built on a Dask parallel computing framework, similar to Minian, enabling scalable processing of large datasets on standard laboratory hardware (e.g. multi-core PCs) without requiring high-performance clusters (***Dong et al.*** (***2022***)). Unlike prior pipelines, MPS enables users to complete every step via an intuitive GUI, created in collaboration with experts in UI/UX design. By combining powerful algorithms with an intuitive interface, MPS aims to lower the barrier for miniscope data analysis revolutionizing the field. This paper describes the design of MPS and demonstrates that it can reliably analyze multi-hour recordings with hundreds of neurons, filling an important need for a scalable and accessible miniscope imaging pipeline. We anticipate that MPS will greatly facilitate the adoption of miniscope technology by the broader community and promote more standardized, reproducible analysis practices.

## Results

### Pipeline Overview & Installation (Figure 1: Step 1)

We developed the Miniscope Processing Suite (MPS), a No-Code GUI-driven pipeline for end-to-end preprocessing, source extraction, and curation of long-duration one-photon calcium imaging that takes an AVI video file from a one-photon imaging experiment and returns a processed temporal and spatial matrix, relevant background components, and a plethora of other intermediate step variables (***Figure 1***A). Unlike CaImAn (***Giovannucci et al.*** (***2019***)), Minian (***Dong et al.*** (***2022***)), or Min1Pipe (***Lu et al.*** (***2018***)), MPS installs via a standalone launcher and runs entirely through a point-and-click interface (https://ariasarch.github.io/MPS_Installer/). Across datasets up to ∼ 2.7 hours (∼ 96,000 frames), MPS completed all stages on a single workstation (Intel^®^ Xeon^®^ w7-3455, Dask memory capped at ∼ 200GB) with a totality of ∼ 7.26TB / ∼ 77 hours of video in a mere 55.6 hours to complete from start to finish. This makes MPS one of the first fully GUI-based pipelines capable of processing tens of terabytes of long-duration miniscope data on a single workstation, highlighting both its accessibility and computational power (***Table 1***).

**Figure 1.**
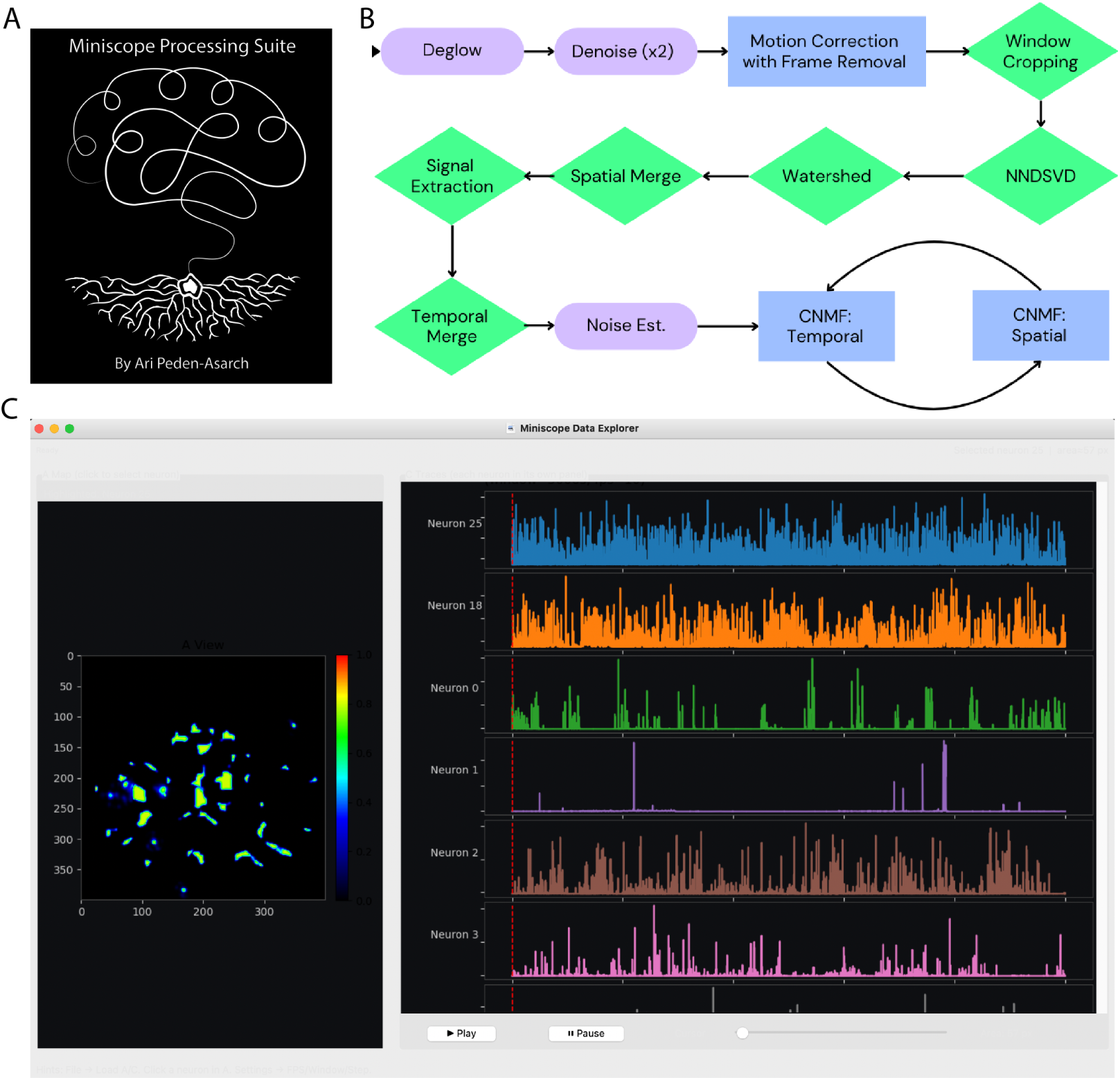
Overview of the Miniscope Processing Suite (MPS): a no-code, end-to-end pipeline for calcium imaging analysis. (A) Logo drawn by Ari Peden-Asarch. (B) Processing pipeline flowchart illustrating the modular architecture of MPS. (C) Interactive data explorer for visualization and quality control of extracted neural signals. Left panel displays spatial footprints of identified neurons color-coded by activity level, with pixel coordinate axes and heatmap scale showing fluorescence intensity. Right panel shows temporal activity traces for selected neurons over 5000 frames.

**Table 1.** MPS algorithmic innovations and empirical performance across 28 recording sessions. Sessions averaged 95,467 frames (86,713–102,874) at 10 Hz, corresponding to 2.4–2.9 hours of recording. Hardware: Intel^®^ Xeon^®^ w7-3455, 8 workers with 2 threads each, Dask memory capped at 200 GB.

| Pipeline Stage | Traditional Approach | MPS Innovation | Median Runtime (min) | % Total |
| --- | --- | --- | --- | --- |
| Preprocessing | FFT-based motion correction | + line-split detection + erroneous frame removal | 37 (IQR: 36–40) | 32% |
| Initialization | 2× least-squares on flattened data [$2 \times M \times T$ in RAM] | NNDSVD + watershed + correlation merging [$0.3 M \times T$] | 26 (IQR: 21–39) | 23% |
| Cell Detection | Direct segmentation | Multi-scale watershed + spatial/temporal merging | 13 (IQR: 8–26) | 11% |
| Validation | Basic filtering | Size/shape/overlap + trace validation + noise estimation | 4 (IQR: 3–6) | 3% |
| CNMF: Temporal | Spatial-first (update all $A$, then all $C$) | **Temporal-first** (filter early for fewer components for spatial) | 7 (IQR: 6–8) | 6% |
| CNMF: Spatial | Global updates on full FOV [$K \times M$ problem] | **KD-tree windowing** [$K_{\text{local}} \times 5\text{--}20\%$ pixels] | 7 (IQR: 5–16) | 6% |
| Export | Standard file I/O | Chunked Zarr, JSON, Pickle, and NPY file export | 7 (IQR: 5–8) | 6% |
| **TOTAL** | Sequential LS-LS-CNMF architecture | Parallel NNDSVD-temporal-spatial architecture | **111 (IQR: 96–137)** | **100%** |

*Processing rate:* MPS achieves 0.7–0.9 min/min of recording. Across 28 sessions totaling ~7.26 TB / ~77 hours of video (up to 2.9 hours per session), MPS completed all stages in 55.6 hours, making it one of the first fully GUI-based pipelines capable of processing tens of terabytes of long-duration miniscope data on a single workstation. Traditional approaches typically would achieve 0.1–0.3 min/min (i.e., up to 10× slower in practice) due to global least-squares initialization, non-windowed spatial updates, and spatial-first ordering.

MPS presents a stepwise GUI (Steps 1-8) with progress bars and easily digestible logs (***Figure 1***B). Purple modules reflect standard preprocessing steps used across existing pipelines: Deglow, Denoise (x2), and Noise Estimation. Green modules represent new innovations introduced in MPS: Window Cropping, NNDSVD initialization, Watershed segmentation, Spatial Merge, Signal Extraction, and Temporal Merge. Blue modules are heavily re-engineered versions of existing algorithms, rewritten to scale to three-hour videos via parallelization: Motion Correction with Frame Removal and iterative CNMF (Spatial/Temporal) updates. Together, this workflow enables MPS to deliver a complete, GUI-driven pipeline for long-duration miniscope imaging. Across 25 installs on macOS and Windows OSes, median setup was 2.5 minutes with 100% first-run success, lowering the barrier for non-programmers. MPS auto-installs its environment and is distributed as a ZIP, DMG, or EXE, making installation seamless across platforms. The Miniscope Data Explorer provides an intuitive environment for quality control after loading spatial footprints (A) and calcium traces (C). Users can interactively inspect each component, visualize its spatial map alongside its temporal activity, and remove low-quality or artifactual cells directly within the interface (***Figure 1***C). A synchronized movie player allows firing activity to be overlaid in real time, enabling visual confirmation of trace fidelity without the need to reconfigure or rerun earlier analysis steps. All edits are non-destructive and can be saved out at any point. The system automatically writes 93 structured files per session, ensuring that data can be re-analyzed globally or revisited on a per-component basis. This design provides both flexibility and reproducibility: investigators can iteratively refine their dataset while maintaining the ability to restore or compare with prior states (***Table 2***).

**Table 2.**
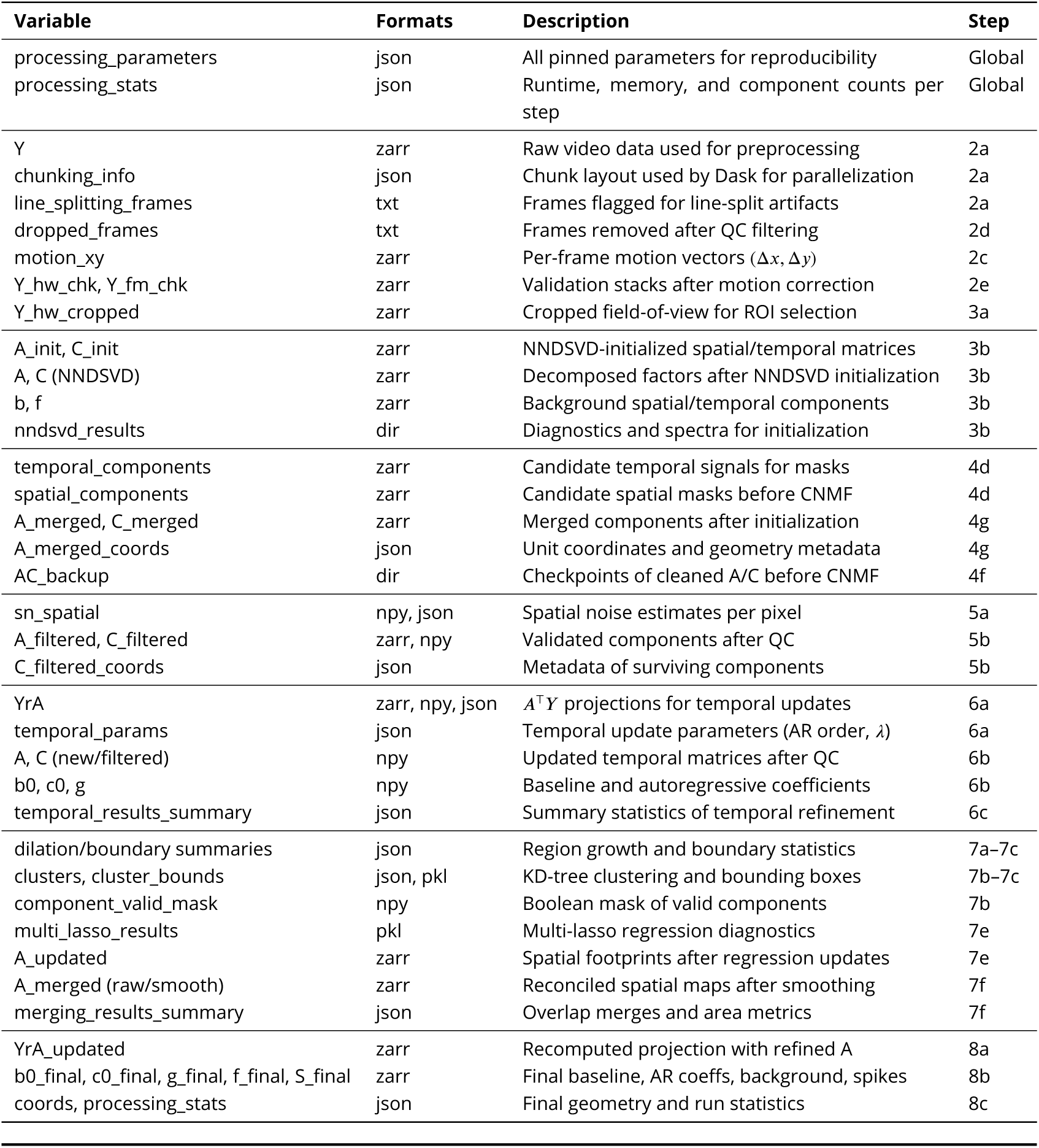
Variables Produced by Each Step of the Miniscope Processing Suite (MPS).

### Video Preprocessing (Figure 2: Steps 2a-2f)

The preprocessing pipeline standardizes video input (***Figure 2***A), subtracts neuropil/background and denoises (***Figure 2***B), performs rigid cross-correlation motion correction (***Figure 2***C), and excludes erroneous frames (line-splits, blur, saturation). On 28 recordings (FOV 600 × 600 pixels, 10 Hz), background suppression reduced the median baseline by 92.59%, and cell-to-surround contrast increased 0.033x (***Figure 2***F). This large reduction is important because neuropil fluorescence dominates raw one-photon miniscope signals; without suppression, somatic activity can be masked by widespread background fluctuations, leading to artificially inflated correlations and reduced sensitivity to single-cell events. Removing this background markedly improves the detectability and interpretability of calcium transients, in line with prior work emphasizing neuropil subtraction in miniscope analysis. Motion correction yielded a residual shift of 3.081 pixels (IQR 2.074 - 4.001) and a long-term drift of ∼0.571 pixels/hr (***Figure 2***G–I). Correcting for motion is critical, as even sub-pixel jitter accumulates over long recordings and can blur spatial footprints, lowering signal-to-noise ratio. Residual drift across hours is especially problematic for neuron registration and stability analysis; mitigating this motion ensures that cell footprints remain well-aligned and neural activity can be tracked reliably across the session.

**Figure 2.**
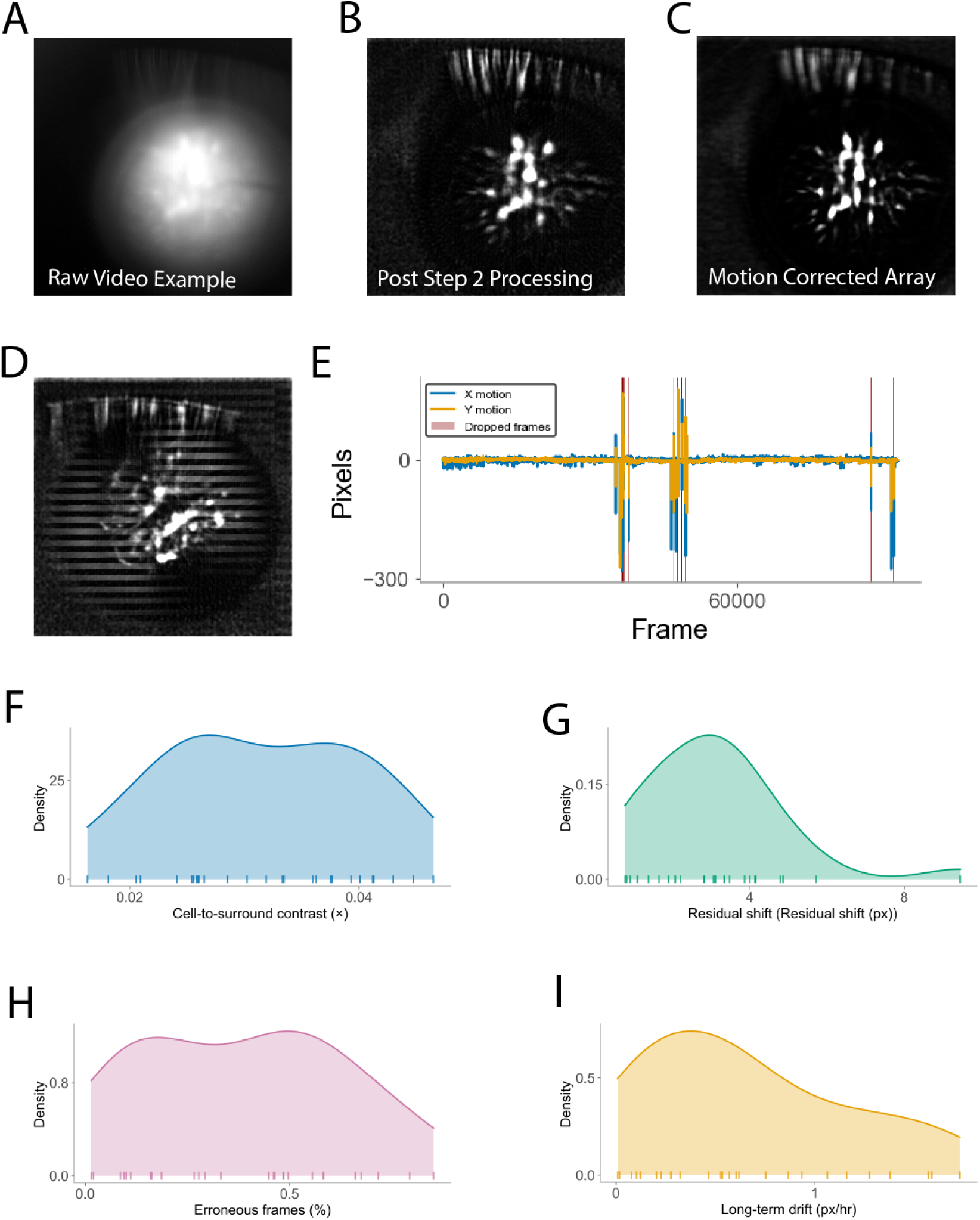
Video preprocessing demonstrates effective background subtraction, motion correction, and quality control. (A) Raw miniscope frame showing diffuse background fluorescence and vignetting that obscures individual cells. (B) Same frame after background subtraction and denoising, revealing individual neuronal somata with improved contrast. (C) Motion-corrected frame showing stable spatial alignment of cellular structures. (D) Concatenated view across multiple frames illustrating line-splitting artifacts (horizontal discontinuities) that are automatically detected and removed. (E) Per-frame motion estimates in X (blue) and Y (yellow) dimensions across the full recording, with erroneous frames highlighted in red bars indicating frames flagged for excessive motion or artifacts (n=518 frames, 0.59% of total of this session). (F) Distribution of cell-to-surround contrast showing improvement after preprocessing (median increase of 0.033×), with individual data points shown as rug plot. (G) Distribution of residual shift after motion correction (median = 3.081 pixels, IQR 2.074–4.001 pixels). (H) Long-term drift quantified as displacement over time (median = 0.571 pixels/hour), demonstrating stability of motion correction across extended recordings. (I) Distribution of erroneous frames as percentage of total recording across all sessions (median = 0.37%), showing automated quality control effectively identifies and removes artifacts without substantial data loss.

Frame quality control flagged on average 0.37% of frames (***Figure 2***I). Although a small proportion, erroneous frames - such as blank captures when the device fails to shut off cleanly, or distorted frames from commutator strain - can strongly skew long recordings (***Figure 2***E). Over three-hour sessions, even rare outliers can bias mean fluorescence, corrupt correlation structure, or disrupt motion correction templates. Automated detection and removal of these frames prevents such disproportionate effects and helps stabilize downstream analyses. Line splitting artifacts, where the top and bottom halves of a frame originate from different time points due to rolling shutter desynchronization, are detected by comparing horizontal discontinuities between adjacent frames; frames exhibiting characteristic mismatched halves (where cross-correlation between frame halves and temporal neighbors exceeds within-frame consistency) are automatically flagged and removed, as demonstrated in our analysis where this method successfully identified and excluded frames with clear temporal tearing artifacts (***Figure 2***D).

### ROI Cropping & Dimensionality Reduction (Figure 3: Steps 3a-3c)

Interactive cropping is applied to remove peripheral or dark regions (***Figure 3***A), which cut the pixel load by 56% (median, 59.0 ± 6.6% mean ± SD across 28 sessions). This reduction decreased run-time by 2.2x (median, 2.52 ± 0.52x mean ± SD) and lowered peak memory usage by 56% (median). Cropping is important because dark borders and unused regions contribute no biological signal but substantially inflate computation time and memory requirements; removing them streamlines analysis without affecting data quality. After cropping, truncated NNDSVD was performed on downsampled frames. With K = 100 components, the decomposition captured 100% of the variance (100.0 ± 0.0% mean ± SD across 28 sessions), with the elbow point typically at K = 3 (range 3-4) (***Figure 3***B–C). Using NNDSVD ensures that the initial spatial and temporal factors are nonnegative and biologically interpretable, while also reducing the number of CNMF iterations compared to random or unconstrained least-squares starts and example of the top three components are displayed for reference (***Figure 3***D). Thus, these reductions enabled full-session analyses to complete 2-3x faster on average while preserving 100% of the explained variance, demonstrating that substantial computational gains can be achieved without any loss in data. Fluorescence photo-bleaching over the 3-hour recording period was minimal, with an average signal decay of 17.4% (***Figure 3***E)). It is important to note that these videos were roughly 60 seconds with a five second off period between them over the 3 hour session to reduce chances of overheating and reduce photobleaching. This gradual decline is automatically corrected by the analysis software’s baseline normalization algorithm, ensuring reliable quantification throughout extended imaging sessions.

**Figure 3.**
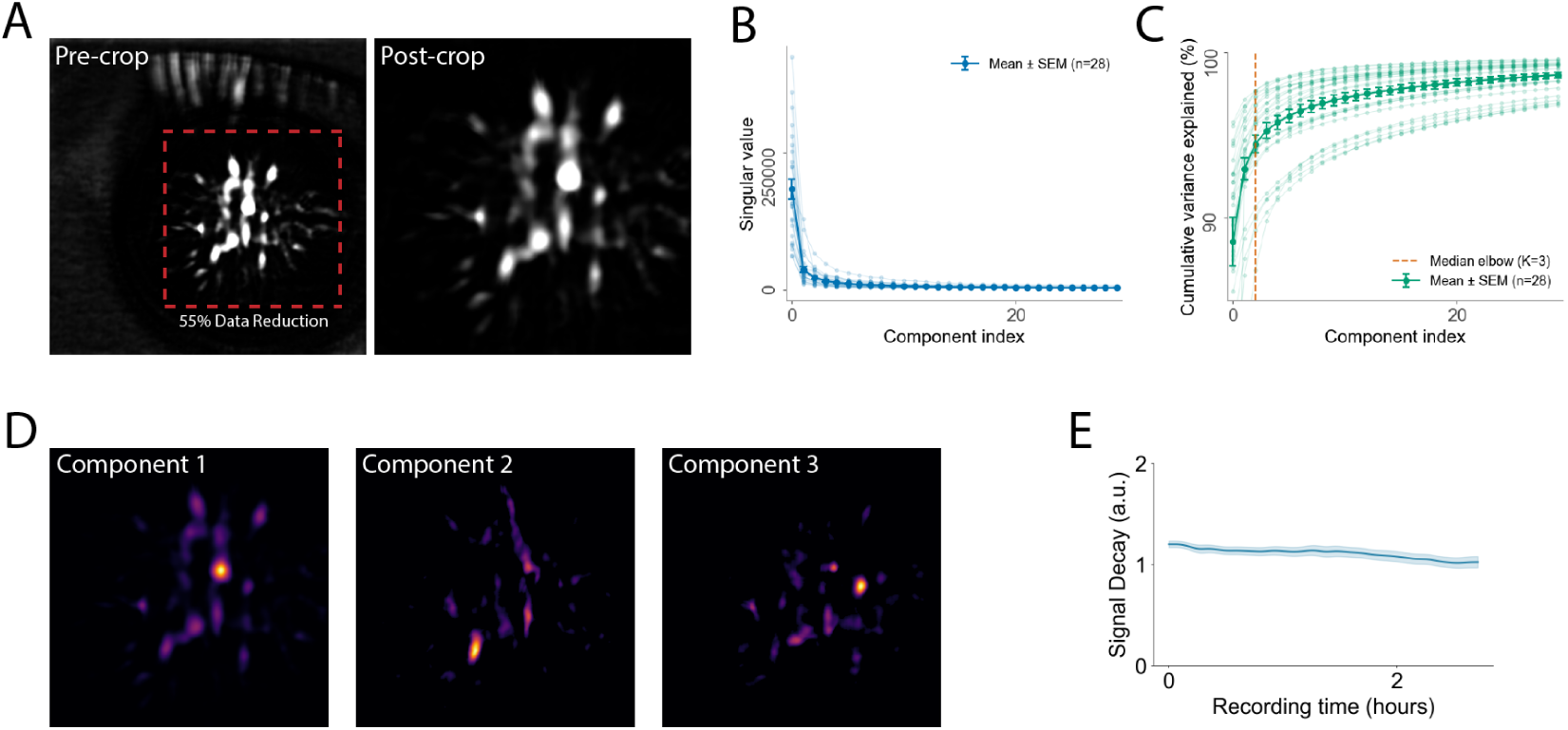
Automated field-of-view cropping improves computational efficiency by reducing pixel load. (A) Pre-crop miniscope frame showing full field of view with vignetting and peripheral regions lacking cellular activity. (B) Post-crop frame demonstrating automated detection and retention of active cellular regions while removing uninformative periphery. (C) Singular value spectrum from NNDSVD decomposition showing rapid decay after initial components, with elbow point indicating optimal dimensionality for downstream processing. Green points indicate selected components capturing majority of signal variance. (D) First three spatial components from NNDSVD decomposition, revealing distinct cellular structures and spatial patterns that serve as initialization for CNMF. Components are ordered by variance explained, with Component 1 capturing the most prominent spatial features and subsequent components revealing progressively finer structures. (E) Fluorescence stability over 3-hour recording period demonstrates minimal photobleaching with mean signal decay of 17.4% with 5s gaps between 60s recordings.

### Cell detection, Initialization, and Pre-CNMF Validation (Figure 4: Steps 4a-5b)

Cells are seeded using local peak detection on maximum and percentile projections, followed by watershed segmentation. Over-segmented masks are then merged by spatial overlap (***Figure 4***A). Initial temporal traces are extracted for each mask and combined with NNDSVD factors to form nonnegative, structured starting matrices for spatial and temporal activity. Components with invalid values are removed, and duplicates are merged if they show both high temporal correlation and spatial proximity (***Figure 4***B) & (***Figure 4***D). In Step 5a, MPS calculates a per-pixel noise map by taking the standard deviation of residuals across time after subtracting a low-rank background estimate. The noise map can be optionally rescaled in regions with strong background activity and smoothed with a Gaussian filter for stability. This provides a component-specific noise level that is later used to scale the sparsity penalty during temporal deconvolution (***Figure 4***F–G). Initialization produced 116 ± 39 candidates per field of view (median area 552 pixels) with a duplicate-merge rate of 98.0% (***Figure 4***B). Across sessions, this pipeline reduced the raw watershed output (5827.2 ± 1359.0 small masks, median size 124.3 ± 20.3 pixels) to 116.0 ± 39.8 curated components (median size 570.7 ± 169.7 pixels) with a 98% duplicate-merge rate, while producing stable noise estimates (overall mean 0.5248 ± 0.0255; range 0.3681-0.9605) (***Figure 4***C) & (***Figure 4***E). Thus, our new approach for seed initialization captures nearly 100% of the variance hours faster than a least squares implementation.

**Figure 4.**
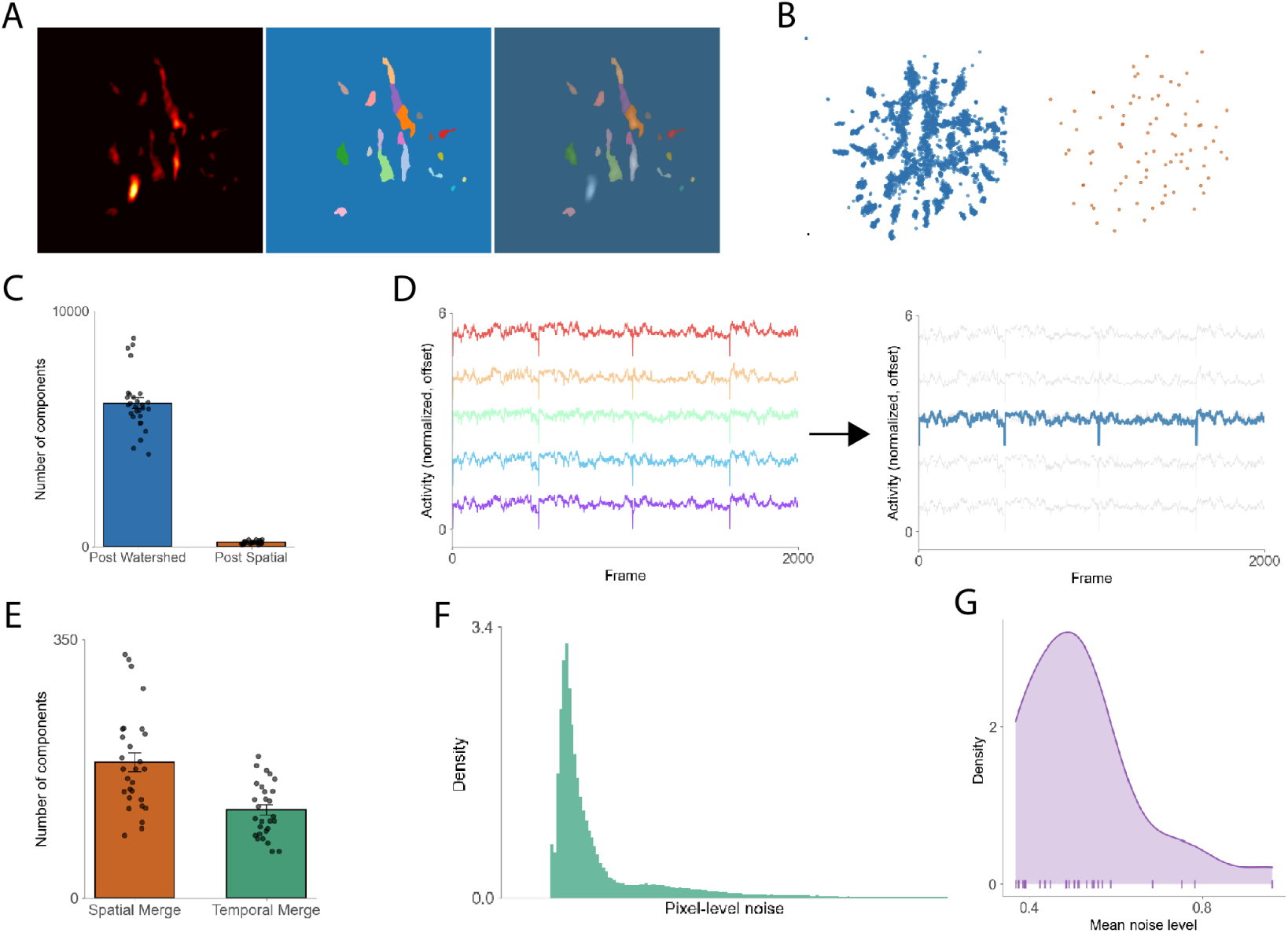
Watershed segmentation and component initialization with spatial merging and validation. (A) Spatial distribution of detected components for Component 1 of an example session showing raw fluorescence image (left), watershed-segmented components with color-coded masks (middle), and merged overlay (right). (B) Scatter plots showing component centroids for all NNDSVD components before (left, high density) and after (right, reduced density) spatial merging. (C) Quantification of component counts across the initialization pipeline, showing reduction from pre-watershed (left) to post-spatial merge (right). Bar plot compares before versus after merging across all sessions with individual data points overlaid. (D) Representative temporal traces from five example components (Comp 2 through Comp 9) showing calcium activity patterns over 2000 frames. Left panel displays raw temporal traces with distinct activity patterns across components; right panel shows activity trace for merged components, demonstrating successful consolidation of correlated signals. (E) Comparison of component counts post-spatial merge (left median = 155.3 ± 53.8) versus post-temporal merge (right, median = 115.3 ± 38.7), demonstrating temporal correlation-based merging further reduces redundant components. (F) Distribution of noise level across all sessions, showing strong peak near real signal and an additional peak towards zero, indicating noise remaining that MPS caught. (G) Distribution of noise estimates caught across all final initialized components after validation, showing median noise level of approximately 0.52.

### Temporal update (Figure 5: Steps 6a-6e)

Temporal activity was updated by projecting the spatial footprints back onto the movie in overlapping time chunks and solving a constrained autoregressive deconvolution. The optimization jointly estimated calcium activity, a nonnegative innovation term with an L1 penalty scaled by each component’s noise level, and baseline terms, subject to nonnegativity and the autoregressive constraint. MPS also suggested decay and sparsity parameters from the noise map and filtered out low-quality components during this step. The residual term (YrA) was centered around zero, indicating that demixing and background subtraction had successfully removed global structure and preserved only noise (***Figure 5***A). On average, components decreased from 114.8 ± 7.6 before filtering to 104.7 ± 6.7 after filtering, corresponding to an overall rejection of 8.8 percent (***Figure 5***E). Median decay time was 1.0 frames with a mean of 1.6 frames, ranging from 0.4 to 20.3 frames across sessions. Signal quality improved markedly, with a mean increase of 55.6 ± 0.8 decibels in signal-to-noise ratio, where signal power was defined as trace variance and noise power was estimated from the median absolute deviation of trace derivative, while signal power was taken as the variance of the trace (***Figure 5***D). Autoregressive decay coefficients yielded median decay of 1.0 frames (100 ms at 10 Hz) and mean of 1.6 frames (160 ms), consistent with the fast kinetics of GCaMP8f indicators (***Zhang et al.*** (***2023***)). Decay times were derived from the autoregressive parameters and summarized with mean, median, and range. It is important to mention that this first pass still yields some noise artifacts, and thus, the subsequent passes described below are needed (***Figure 5***B–C).

**Figure 5.**
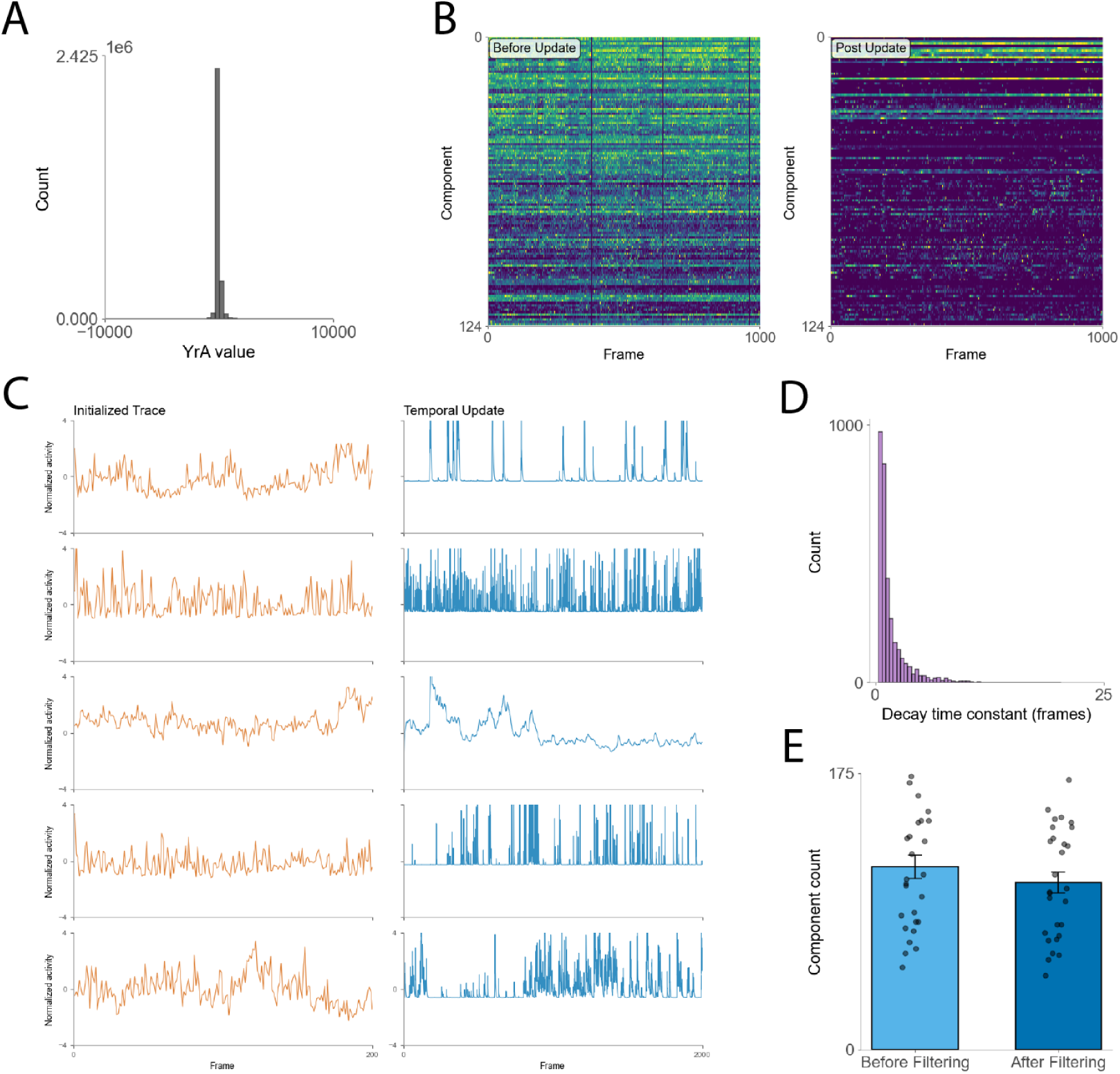
Temporal activity update through autoregressive deconvolution with quality control filtering. (A) Distribution of YrA values (residual after projecting spatial components onto the movie) centered near zero, indicating successful demixing and background subtraction. (B) Heatmaps of temporal activity for all components for a single session before (left) and after (right) the temporal update. Each row is one component. Color intensity represents normalized calcium activity across frames. (C) Representative example traces showing the initialzied trace (left, 5 examples) and trace after the temporal update (right, 5 examples labeled). (D) Distribution of estimated decay time constants across all components, showing median decay of 1.0 frames (mean = 1.6 frames, range 0.4-20.3 frames). (E) Component retention across temporal filtering, comparing counts before (left, median = 114.8 ± 7.6) versus after (right, median = 104.7 ± 6.7) quality control, representing 8.8% rejection rate. Signal-to-noise ratio improved by mean of 55.6 ± 0.8 dB across retained components.

### Spatial update (Figure 6: Steps 7a-7f)

Spatial footprints were refined using localized, neighbor-aware nonnegative regression. Each component’s region was dilated, local residuals were recomputed with neighbor subtraction, and footprints were updated under nonnegativity and locality constraints (***Figure 6***A). Heuristic checks flagged components with implausible sizes, updated masks were adopted, and spatial duplicates were merged. Component counts dropped substantially, from 104.7 ± 6.7 before refinement to 51.6 ± 4.1 after, a reduction of 50.8 percent (***Figure 6***B). Nearest-neighbor statistics showed increased spacing, with mean centroid distance rising from 19.6 ± 10.5 to 27.0 ± 14.5 pixels and median distance increasing from 19.0 to 26.2 pixels (***Figure 6***C). After a single update on the spatial dimension, the location of the cells can now easily be discerned.

**Figure 6.**
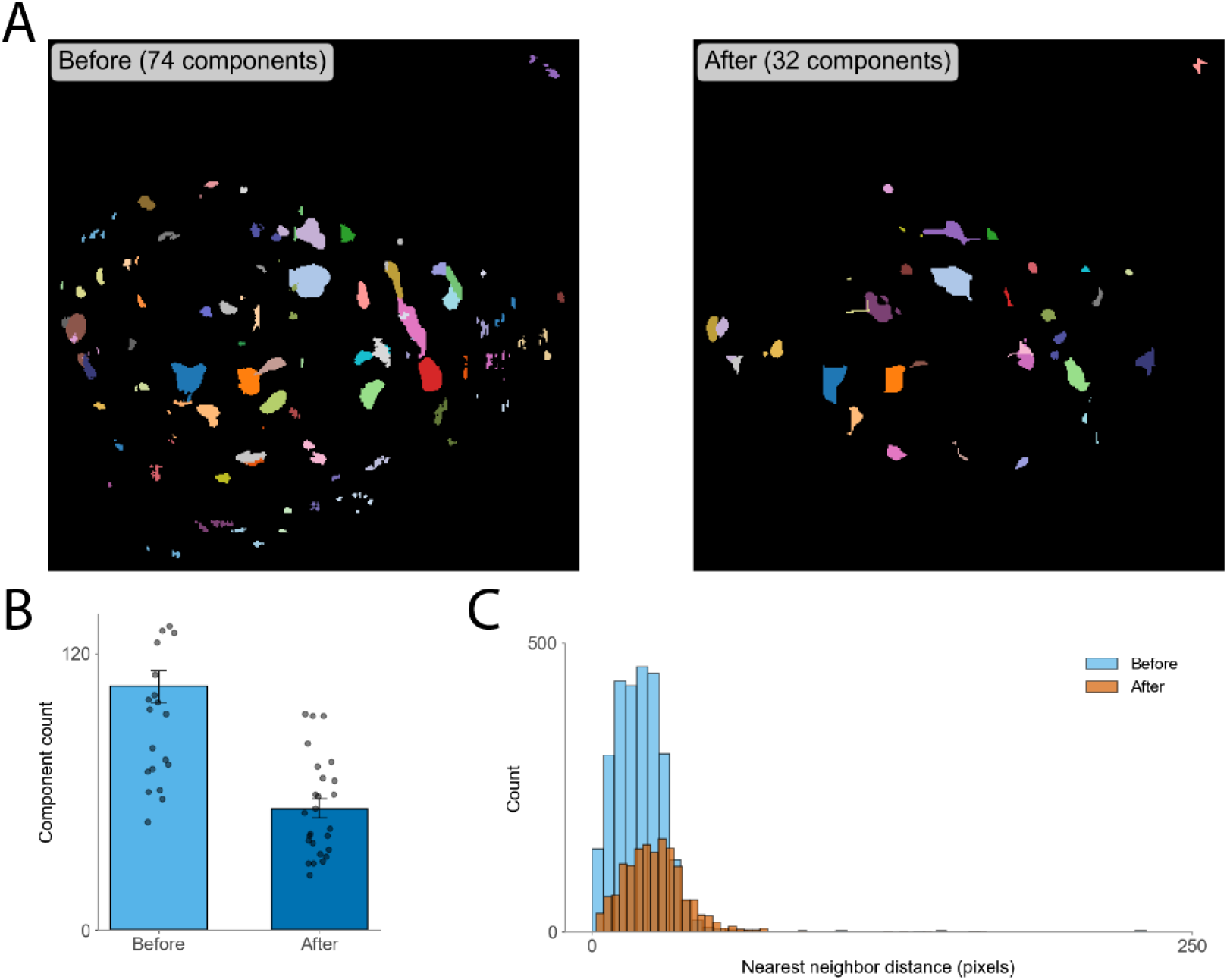
Spatial footprint refinement through localized nonnegative regression with neighbor-aware updates. (A) Spatial distribution of components before (left) and after (right) spatial refinement. Each colored region represents refined spatial footprint after dilation-bounded regression and neighbor subtraction. Post-refinement components show improved spatial localization and reduced overlap. (B) Component count reduction across sessions from pre-spatial update (median = 104.7 ± 6.7) to post-spatial update (median = 51.6 ± 4.1), representing 50.8% reduction through merging of spatial duplicates and removal of implausible footprints. (C) Distribution of nearest-neighbor distances between refined component centroids, showing shift toward larger separations with peak around 100-150 pixels and histogram counts up to 400 components.

### Final Temporal Update (Figure 7: Steps 8a-8c)

The final temporal refinement was performed after spatial components had been merged and residuals refreshed. This two-stage temporal refinement strategy (updating traces both before and after spatial refinement) follows the iterative optimization protocol established in CNMF-E, where one additional matrix update is typically performed following component interventions to ensure convergence (***Zhou et al.*** (***2018***); ***Giovannucci et al.*** (***2019***)). As in the earlier update, the spatial footprints were projected back onto the movie in overlapping time chunks, and a constrained autoregressive deconvolution was solved to jointly estimate calcium activity, a nonnegative innovation term with sparsity regularization, and baseline parameters. At this stage, the optimization incorporated the merged spatial set and the updated residuals, providing a definitive estimate of temporal activity. Low-quality traces were pruned, and background terms could be optionally modeled alongside component activity. The residual term (YrA) was even more centered around zero, indicating that demixing and background subtraction had better removed global structure and preserved only noise when compared to the first temporal pass (***Figure 7***A) This final pass substantially reduced the number of surviving components, from 117.1 ± 7.7 before filtering to 52.6 ± 4.2 after filtering, corresponding to a 55.1% rejection rate (***Figure 7***E). Median decay time remained 1.0 frames with a mean of 1.6 frames, spanning 0.4–20.3 frames across sessions (***Figure 7***D). Signal quality was further improved, with a mean increase of 47.8 ± 1.2 dB in signal-to-noise ratio. These results established the final set of temporal traces used in downstream analyses (***Figure 7***B–C). Additionally, if one wishes to continue to perform temporal or spatial updates, they are more than able to through MPS, but not required. See Supplementary Video 1 for an example of the final Miniscope Processing Suite output.

**Figure 7.**
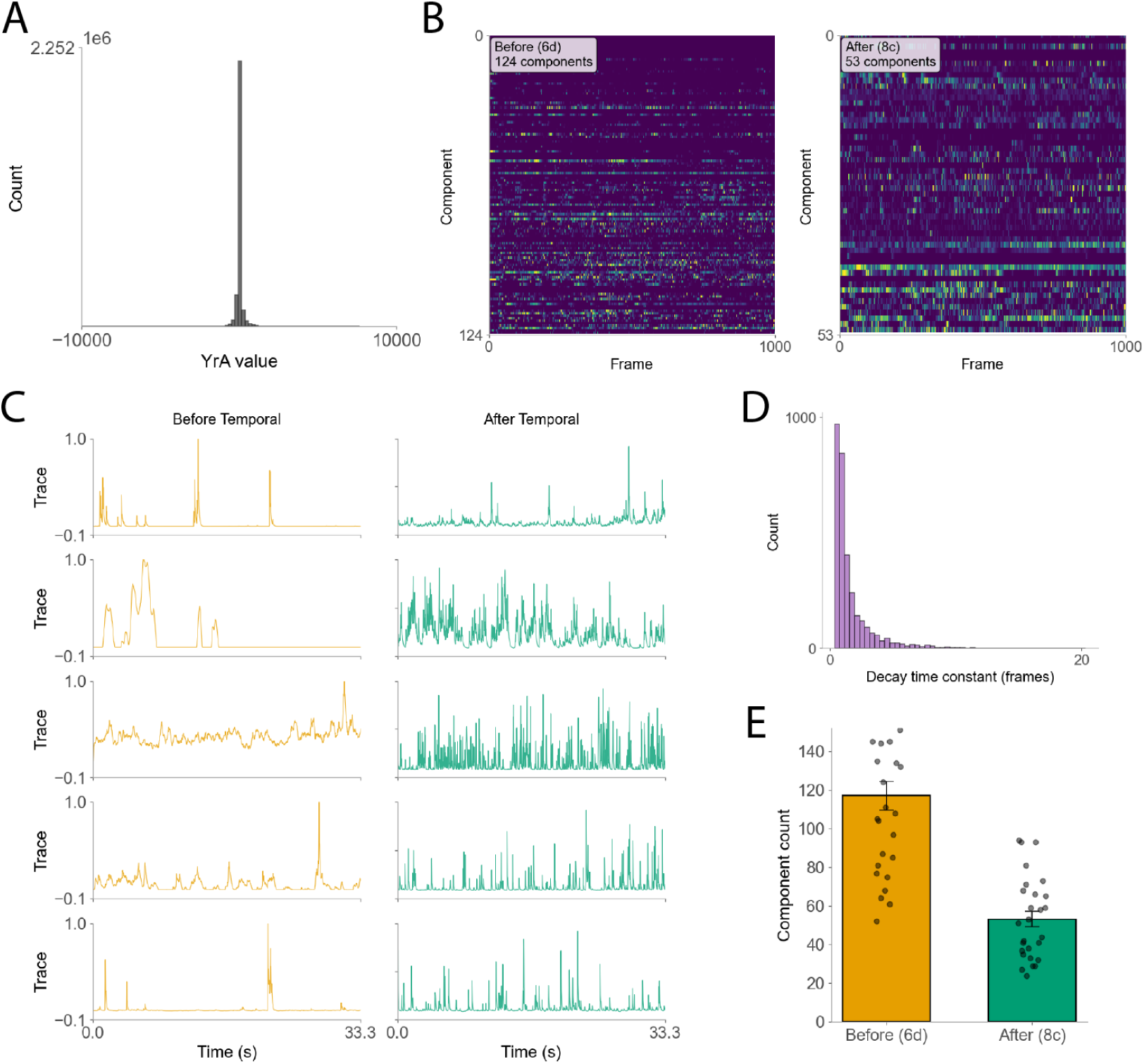
Final temporal refinement with updated spatial footprints produces high-quality curated neural components. (A) Distribution of YrA values (residual after projecting refined spatial components onto the movie) centered near zero, indicating successful demixing and background subtraction with improved spatial footprints compared to first temporal pass. (B) Heatmaps of temporal activity for all components for a single session before (left) and after (right) the final temporal update. Each row is one component. Color intensity represents normalized calcium activity across frames. (C) Representative example traces showing post-first-pass temporal update traces (left, 5 examples) and denoised calcium traces after the second temporal refinement (right, 5 examples). (D) Distribution of estimated decay time constants across all final components, showing median decay of 1.0 frames (mean = 1.6 frames, range 0.4-20.3 frames), consistent with first temporal pass. (E) Component retention across final temporal filtering, comparing counts before (left, yellow/gold bar, median = 117.1 ± 7.7) versus after (right, teal bar, median = 52.6 ± 4.2) quality control, representing 55.1% rejection rate.

### Summary

MPS combines (i) interactive cropping to avoid analyzing the uninformative parts of the video, (ii) NNDSVD to provide nonnegative, interpretable initial factors without unconstrained least-squares, and (iii) parallel, out-of-core temporal (AR(2), CVXPY) and spatial updates, producing accurate, scalable analyses of multi-hour recordings on standard lab hardware. The integrated Data Explorer closes the loop with transparent manual QC, supporting rigorous and reproducible workflows.

## Discussion

### Accessibility

A primary advantage of MPS is its accessibility to users of varying computational backgrounds. Unlike prior calcium imaging pipelines that require deep programming knowledge or command-line operation (e.g. CaImAn, MIN1PIPE, and CNMF-E in Python/MATLAB, or Minian via Jupyter notebooks), MPS provides a fully graphical user interface that guides the experimenter through each analysis stage. All major functions can be executed with menu selections and button clicks, with no need to modify code. All code is open-source, with detailed comments to facilitate transparency and ease of use. This GUI/no-code design dramatically lowers the barrier to entry. Users who are unfamiliar or intimidated with coding can now easily access miniscope processing technology through MPS, democratizing the analysis of multi-hour long miniscope data (***Dong et al.*** (***2022***)). The software includes built-in tooltips, tutorials, default parameter presets, full automation of processing, and a data explorer platform further simplifying usage for beginners. Thus, what was once a scripting-intensive workflow was transformed into an interactive application accessible to the broader scientific community via MPS. This improved usability addresses a key limitation of earlier pipelines, wherein hard-coded parameters and lack of user guidance often made it difficult to trust and validate results without extensive tinkering. By transforming advanced calcium imaging analysis into a point-and-click process, MPS allows researchers to focus on experimental questions rather than computational setup and system management. Additionally, these results can be easily repeated or shared since the GUI enforces a consistent sequence of steps and settings, in contrast to ad-hoc custom scripts. Overall, the accessibility of MPS is poised to empower many labs to adopt miniscope imaging that previously found the data analysis too daunting (***Freeman*** (***2015***); ***White et al.*** (***2020***); ***Aharoni et al.*** (***2019***)).

### Scalability for Longer Recordings

Beyond usability, MPS achieves scalability rarely attainable in existing miniscope analysis software. Benchmarking across sessions up to three hours demonstrated stable performance and predictable memory behavior even on constrained workstations. By combining out-of-core Dask execution, blockwise CNMF updates, and efficient chunking of intermediate arrays, MPS processes continuous videos exceeding 95,000 frames (∼ 2.7 hours) without requiring manual chunking or external cluster resources. This is a significant improvement over prior implementations that saturate memory or fail during motion correction. Runtime scales linearly with dataset duration, and the memory footprint remains bounded by user-defined caps. Even when capped at 200 GB of memory, the full pipeline completed processing of 7.26 TB of miniscope data (total 77 hours of raw video) within 55.6 hours on a single Xeon workstation. This scalability ensures that experiments demanding high temporal coverage or long-term tracking can now be analyzed on ordinary lab hardware, promoting reproducibility and widespread adoption.

### Extensibility to New Modalities

MPS was intentionally designed with modularity, allowing extensions beyond calcium imaging. Each analysis step is encapsulated as a standalone module within the pipeline framework, with standardized data interfaces that make it straightforward to integrate new frontends or replace existing algorithms. For example, future versions could incorporate voltage imaging, fiber photometry, or photometric motion tracking modules. Similarly, deep-learning models for ROI detection or denoising can be plugged into the pipeline through the model integration interface, which already supports loading pre-trained models from external frameworks such as PyTorch or TensorFlow. This modularity ensures that MPS can evolve alongside new imaging modalities and community-driven innovations while retaining the same no-code workflow.

### Limitations

While MPS expands accessibility and scalability, several limitations remain. The current design focuses on preprocessing and CNMF-based source extraction; online or real-time decoding is not yet implemented. Additionally, although Dask parallelization enables large-scale computation on a single workstation, true distributed or cluster-scale processing remains future work. GPU acceleration, while feasible for some submodules, is not yet uniformly implemented due to hardware-specific dependencies. Future updates may include optional CUDA-accelerated paths and additional preprocessing algorithms to further improve performance. Another limitation is that long recordings with highly dynamic fluorescence or photobleaching may require customized baseline correction routines not yet part of the GUI presets. Nevertheless, these limitations represent tractable engineering improvements rather than conceptual barriers.

### Community Impact

The primary impact of MPS lies in democratizing advanced calcium imaging analysis. By enabling non-programmers to perform analyses that were previously accessible only through scripting, MPS lowers the technical threshold for neuroscience research. This in turn fosters inclusivity and reproducibility, aligning with broader open-science principles. The code base, released under an open-source license, encourages transparency and collaboration, while the GUI ensures consistent step ordering and parameterization across users. These design choices promote a unified standard for miniscope data analysis that can serve as a foundation for future benchmarking and cross-lab validation studies. Ultimately, MPS empowers the neuroscience community to shift focus from technical barriers to scientific discovery, accelerating progress in systems neuroscience.

## Methods and Materials

### Step 0: Download

MPS provides comprehensive installation support across multiple platforms, ensuring broad accessibility for the neuroscience research community. The software is distributed through a dedicated installation website (https://ariasarch.github.io/MPS_Installer/) that offers platform-specific installers optimized for different operating systems. For macOS users, MPS is available in both ZIP and DMG formats, supporting macOS 12+ on both Apple Silicon and Intel architectures. The application is signed and notarized, ensuring security compliance and seamless installation without security warnings. Windows users can access a dedicated EXE installer compatible with Windows 10/11 (64-bit systems), requiring approximately ∼1GB of disk space during the initial setup process.

The installation process is fully automated and consists of four key steps: (1) downloading the MPS payload from the GitHub repository, (2) creating appropriate shortcuts on Windows or a self-contained application on macOS, (3) establishing the Python environment during the first launch, and (4) enabling instant subsequent launches. This streamlined approach minimizes technical barriers for researchers while maintaining robust functionality. System requirements are modest: ∼16GB RAM minimum (∼32GB recommended), stable internet connectivity for initial setup, and the aforementioned disk space allocation. The software architecture prioritizes accessibility through its intuitive graphical interface while maintaining the computational power necessary for complex calcium imaging analysis. Support infrastructure includes comprehensive documentation, an active issue tracking system for bug reports and feature requests, and community discussion forums (https://github.com/ariasarch/MPS_1.0.0/discussions/). The complete source code is available on GitHub, enabling customization for specialized laboratory requirements and fostering collaborative development within the research community (https://github.com/ariasarch/MPS_1.0.0/).

### Step 1: Graphical User Interface, Environment Provisioning, and Automation

MPS is implemented in Python (tested on Python 3.8) and distributed as a standalone, point-and-click application. To minimize package-management burden, the distribution includes a one-click launcher that triggers when the icon is clicked and provisions the Python runtime and pinned dependencies before starting the GUI; no user interaction with package managers such as conda, mamba, and pip is required. The main application imports all step definitions at launch and exposes them through a centralized controller, ensuring that the GUI can orchestrate modules consistently across runs. Additionally, all analysis parameters are centrally managed and saved to a single JSON configuration, enabling exact reproducibility and seamless resumption. The configuration captures both global settings (e.g., worker count, memory limits, and dataset paths) and per-step options (e.g., segmentation thresholds, NNDSVD rank, deconvolution hyperparameters). Representative keys are included and persisted alongside the dataset metadata for easy referencing.

Upon launch, the GUI presents a stepwise workflow with a status bar and progress indicator. Users can navigate the pipeline incrementally (Previous/Next) or operate it in fully automated mode. Moreover, a background process checks for current processing to protect the user from moving from step to step prematurely and risking a loss of data. Automation controls are exposed via the Automation menu allowing hands-off execution from any stage. An autorun indicator in the navigation bar reflects the current automation state, and the default delay between steps is user-settable. Users are advised to gain familiarity with the software and its parameters before enabling automation to ensure appropriate configuration. Parallel execution is handled automatically by the system; however, advanced users can access the Dask Dashboard via the ‘Open Dask Dashboard’ option, which provides a live link for monitoring task progress, memory usage, and worker activity. Moreover, for iterative work, a Load Previous Data dialog assists in restoring prior sessions detecting available cache artifacts (e.g., .zarr, .npy, .json), lets users select which to load, and can mark steps “completed through” a chosen stage so that processing resumes exactly where intended in case of an unexpected session termination. Parameters are auto-saved on exit to prevent configuration drift between runs as well.

Together, these design choices such as, environment provisioning via a one-click launcher, centralized JSON parameterization, menu-level automation toggles, optional notifications, and live dashboard access lower the operational complexity of running a full CNMF pipeline while preserving rigorous control over analysis settings.

### Step 2: Data Preprocessing

Step 2 prepares raw miniscope video data through a series of cleaning operations to enhance signal quality and reduce artifacts prior to source extraction. This preprocessing pipeline is subdivided into Steps 2a–2f, each described below. For all subsequent cost analysis, let *T* be the number of frames in the recording and *M* the number of pixels per frame.

### Step 2a: Video Loading and Format Standardization

The pipeline begins with loading miniscope videos via FFmpeg and materializing them as an xarray backed by Dask (lazy, chunked array), enabling out-of-core processing instead of holding full movies in RAM. The loader uses a regex file-pattern to select inputs and, in the current build, defaults to the AVI video format commonly saved by the Miniscope V4. For each file, MPS ffprobes the stream and record width, height, and frame count and log the file path and size for traceability. Frames are read through a delayed FFmpeg decode and wrapped into a Dask array of dtype uint8 with dimensions [frame, height, width]. The resulting array can be optionally cached on disk as a chunked store for reuse in later steps. An optional automated screen flags frames consistent with the “line-splitting” artifact. In the current implementation, the mean intensity is computed for every frame in the leftmost 20 columns and mark frames whose left-edge mean exceeds a robust threshold (global mean + 2× the Standard Deviation across frames):

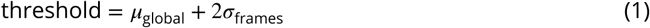

Indices of flagged frames are saved, logged, and removed from the xarray before downstream processing. This straightforward yet effective safeguard is absent from existing pipelines, underscoring MPS’s emphasis on practical quality control. This built-in QC prevents obviously corrupted frames from propagating into motion correction and demixing. Computationally, byte I/O and decoding scale linearly with dataset size, and thus, the total cost is *O*(*M* × *T*). The line-split screen adds a small constant-factor pass over a fixed-width strip (∼ 20× height per frame).

### Step 2b: Background Subtraction and Denoising

Miniscope recordings often suffer from uneven illumination and pixel noise. MPS addresses these by first subtracting a static background image from the video. Concretely, a per-pixel low-percentile projection (default: 1–5% or the minimum over a representative subset of frames) is computed to estimate the persistent background fluorescence (vignetting and baseline autofluorescence) under the assumption that each pixel intermittently approaches baseline during the session (***Giovannucci et al.*** (***2019***)):

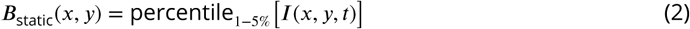

This background image is subtracted from every frame, removing constant offsets while preserving relative calcium fluctuations:

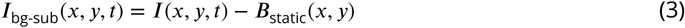

Users can choose among median filtering (robust to salt-and-pepper noise), Gaussian (smooths gradual pixel-level variation), bilateral (preserves edges while denoising), or anisotropic diffusion (iterative edge-preserving smoothing for stronger noise regimes), giving flexibility to tailor denoising to different noise profiles and experimental conditions. The most common option recommended is a frame-wise median filter (3×3 kernel) suppresses high-frequency salt-and-pepper noise without blurring soma edges (***Giovannucci et al.*** (***2019***)):

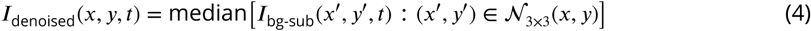

Finally, morphological opening is performed with a disk-shaped structuring element with the radius matched to typical soma radius; e.g. 10 pixels to estimate and remove diffuse neuropil/background glow (***Lu et al.*** (***2018***)):

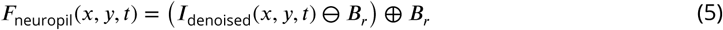

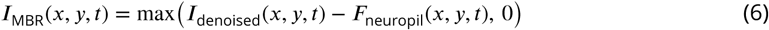

This “rolling-ball” equivalent operation has a long history in biomedical image processing and was introduced to miniscope pipelines in MIN1PIPE (***Sternberg*** (***1983***); ***Lu et al.*** (***2018***)). Each filter operates independently per frame, so the total cost is *O*(*M* × *T*) and parallelizes across CPU workers. At the end of Step 2b, the data now has a near-zero baseline and reduced high-frequency noise, improving robustness of downstream motion correction and cell detection.

### Step 2c: Motion Correction by Cross-Correlation

Head-mounted miniscopes often exhibit motion artifacts due to physical movement of the brain in the skull. In Step 2c, MPS performs rigid motion correction on the preprocessed video using template matching with normalized cross-correlation computed in the Fourier domain (***Lewis*** (***1995***)). A reference image is defined (by default, the mean projection after background subtraction; alternatively, a max projection can be used for higher contrast). For each frame, the 2D normalized cross-correlation is evaluated with the reference to obtain the translation (Δ*x*, Δ*y*) that maximizes similarity:

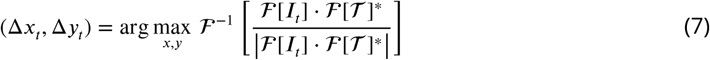

To handle large displacements efficiently, a coarse-to-fine strategy is used: a downsampled (pyramidal) pass provides coarse shifts, followed by a high-resolution pass to refine alignment. Sub-pixel accuracy is achieved by fitting the correlation peak via an upsampled DFT around the maximum (***Guizar-Sicairos et al.*** (***2008***)). Frames in which peak correlation falls below a user-set threshold (blur, occlusion) are flagged as unreliable and excluded from downstream estimation. The FFT-based correlation for one frame costs *O*(*M* log *M*); over *T* frames the total is *O*(*T* × *M* log *M*). Parallelization across frames (CPU workers) is used to accelerate this step. The output is a per-frame shift vector applied in Step 2e to remap the movie into a stable coordinate frame. For context, non-rigid/piecewise-rigid alternatives (e.g., NoRMCorre) are common in calcium imaging (***Pnevmatikakis and Giovannucci*** (***2017***)); here a rigid variant is adopted, which is sufficient after denoising and background suppression in Step 2b.

### Step 2d: Detection of Erroneous Frames

After motion estimation, MPS automatically screens for frames with excessive inter-frame motion and optionally removes them. The detector consumes the per-frame x/y shift estimates produced during motion correction and flags frames in which shift magnitude is an outlier relative to the run’s typical motion:

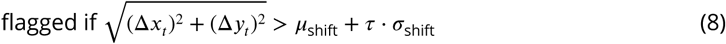

Concretely, for each motion dimension present (vertical/height, horizontal/width), the mean and standard deviation of the estimated shifts is computed and mark any frame in which shift exceeds a user-tunable threshold factor (default *τ* = 5.0) times the standard deviation from the mean. When the “Drop Erroneous Frames” option is enabled (default: on), the flagged frames are removed in tandem from both the motion dataset and the reference video array carried forward from Step 2b; the arrays are then rechunked to keep frame-chunk sizes well-formed for downstream Dask operations. MPS writes two audit files: a per-step list of dropped indices (dropped_frames.txt) and a combined list (all_removed_frames.txt) that merges Step 2a line-split removals with Step 2d motion outliers, adjusting indices back to the original frame numbering for reproducibility. This screening is linear in the number of frames i.e., *O*(*T*) with a small constant factor. Because the underlying arrays are Dask-backed, the work is executed out-of-core and can leverage available workers allowing for multi-hour recordings be sanitized without ever loading the full movie into RAM, avoiding scheduler stalls from oversized task graphs, and preventing a handful of extreme-motion outliers from corrupting templates and biasing downstream CNMF.

### Step 2e–2f: Frame Transformation and Remapping and Post-Validation of Preprocessing

Once the optimal shifts are determined, MPS applies a per-frame rigid warp to generate a stabilized movie. Each frame is translated by its (Δ*x*, Δ*y*) estimate and resampled with sub-pixel interpolation. The GUI exposes these options directly and saves the corrected data as a Zarr dataset, with optional derivatives chunked for time-series or spatial access to support downstream steps. The operation is linear in data size (*O*(*M* × *T*)) and parallelized across frames using Dask. Sub-pixel remapping follows standard image-registration practice (***Thévenaz et al.*** (***1998***); ***Guizar-Sicairos et al.*** (***2008***)). After Step 2e, MPS provides a validation panel that runs lightweight checks on the motion-corrected video array before advancing to Step 3. Users can set a sample size for analysis (default 1000 frames; 0 means all frames) and enable three checks: (i) frame-intensity statistics, (ii) NaN/Inf detection, and (iii) an optional chunking comparison that benchmarks two storage layouts carried forward from Step 2e (frame-chunked vs spatial-chunked). These checks operate on the already-produced Step 2e arrays and execute via Dask out-of-core computation, and thus, runtime scales approximately linearly with the number of frames evaluated (*O*(*T*) for the chosen sample).

### Step 3: Field-of-View Cropping and Dimensionality Reduction

To reduce computational load and focus analysis on relevant regions, MPS applies interactive cropping of the field of view followed by truncated NNDSVD for dimensionality reduction. This step removes non-informative pixels, accelerates downstream processing, and provides biologically interpretable initialization factors for CNMF

### Step 3a: ROI Cropping

Miniscope videos often contain dark or irrelevant regions e.g., the circular vignette periphery or tissue outside the target area (***Leong et al.*** (***2003***)). MPS provides an interactive cropping tool that lets the user define a focused field of view via a radius slider (default 75% of the smaller image dimension) and X/Y offsets, with “Preview Crop” and “Apply Crop” actions and a “Use Full Frame” reset. The preview overlays the proposed crop on a mean-intensity image; applying the crop saves chunked Zarr arrays for downstream steps and logs progress in the GUI. All pixels outside the selected region are discarded for subsequent analysis. This simple operation typically reduces pixel count by ∼ 70% for a 0.6mm GRIN Lens (Inscopix-distributed GRINTECH optics), yielding a comparable speedup in later steps whose cost scales with the number of pixels *M* (e.g., a 70% reduction in *M* gives ∼ 3.3× faster computation). Implementation is a per-frame mask-and-trim (rectangular crop aligned to the selected center), with total complexity *O*(*M* × *T*) and trivial parallelism across frames. After cropping, all frames are truncated to the crop’s bounding box, and *M* thereafter denotes the pixel count with this new ROI.

### Step 3b–3c: Non Negative Double Singular Value Decomposition (NNDSVD) for Dimensionality Reduction

To obtain a compact, nonnegative initialization for CNMF, the cropped, motion-corrected video dims is reshaped: [frame, height, width] to a 2-D matrix [frames, pixels] by stacking the spatial axes, then shift the data to nonnegativity by subtracting the global minimum and adding a small epsilon. An in-memory randomized SVD with user-selectable parameters (#components, default 100; power iterations, default 5). This yields *U*, *S*, and *V* ^⊤^ where *U* ∈ ℝ^*T* ×*K*^, *S* ∈ ℝ^*K*^, and *V* ^⊤^ ∈ ℝ^*K*×*M*^ is then attempted:

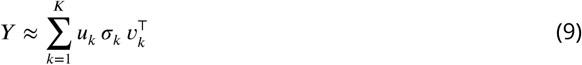

From these, an NNDSVD is constructed by taking thresholded positive/negative parts of each singular triplet and (optionally) applying a spatial localization score to prefer concentrated spatial maps:

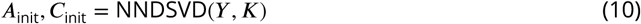

The first component is reserved as a background factor; the remaining components seed potential neurons. The resulting initial factors are written to disk as *A*_init_ [component, height, width], *C*_init_ [frame, component], and background components in space and time (*b* and *f* respectively) for the spatial and time background. (***Boutsidis and Gallopoulos*** (***2008***); ***Halko et al.*** (***2011***)) Additionally, this step has a near-linear pass over the data with additional *O*((*M* + *T*)*K*^2^) terms, and thus, run-time scales with *K* components, frames, and pixels. After initialization, Step 3c provides a brief diagnostics and QA for the NNDSVD results. The GUI renders: (i) singular value spectrum and cumulative variance explained; (ii) a sparsity comparison (non-zeros in the raw SVD vs. the NNDSVD); (iii) an overlap map showing how many components cover each pixel; (iv) a background component panel (spatial map, temporal trace, histogram, and power spectrum); (v) per-component size distributions, relative singular values, max spatial intensities, and temporal variability; (vi) a component details view (spatial map, temporal trace, spatial profiles, and summary stats). Users can also export a CSV of component metrics for records/review.

### Step 4: Initial Cell Detection and Component Initialization

Step 4 constitutes the initialization of potential cell components (spatial footprints and temporal traces) from the preprocessed video. MPS employs a two-pronged strategy for initialization: (1) image segmentation to directly detect cell locations and (2) matrix factorization-based initialization to decompose the video into the relevant components. By uniting direct image segmentation with matrix factorization, MPS introduces a previously unexplored initialization strategy that delivers an exceptionally robust and biologically faithful foundation for CNMF refinement.

### Step 4a–4b: Peak Detection via Watershed Segmentation

To initialize potential cell footprints, MPS applies watershed-based segmentation to a representative projection image. By default a maximum-intensity projection is computed from the motion-corrected data so that active neurons appear as bright peaks; the image is lightly smoothed with a Gaussian kernel (multi-scale search, *σ* ∈ [0, 2] pixels) and local maxima are detected subject to a minimum inter-peak distance (typically 10–15 pixels) and a relative intensity threshold (e.g., > 10% above local background). These peaks serve as markers for a marker-controlled watershed on the inverted projection, partitioning the image into candidate regions whose boundaries follow intensity saddles around each peak (***Vincent and Soille*** (***1991***); ***Lu et al.*** (***2018***); ***van der Walt et al.*** (***2014***)). Parameters are biased toward slight over-segmentation at this stage (favoring sensitivity to dim or irregular somata) since redundant or irrelevant fragments are merged or dropped respectively downstream. The worst-case time complexity of the watershed transform is *O*(*M* log *M*) where *M* is the number of frames, but in practice is near-linear in the number of pixels for 2D images of this size. In the GUI, Step 4a performs a small grid search over min_distance, threshold_rel, and *σ* on a subsample to propose sensible defaults; results (including the best triplet and summary plots) are written to state and auto-loaded by Step 4b, which then applies marker-based segmentation across components with optional minimum-region-size filtering and parallel execution. These controls are exposed directly in the Step 4a/4b panels (e.g., Run Parameter Search, Run Segmentation, Skip Background, Use Parallel Processing) and their choices are persisted to the session configuration for exact reproducibility.

### Step 4c: Spatial Merging of Candidate Segments

MPS merges spatially proximate candidate segments from Step 4b using simple, size-aware centroid heuristics since the watershed segmentation usually produced a drastic overestimation of neural components. The step first loads the component list produced by Step 4b and optionally filters out very small segments (default min size = 9 pixels). The user controls four parameters in the GUI: Distance threshold (default 25 pixels), Size-ratio threshold (max larger/smaller, default 5.0), Minimum size (default 9 pixels), and Maximum merged size (default 5000 pixels). Merging proceeds greedily as follows: for each seed component not yet processed, MPS initializes a group and scans subsequent components where a candidate is added to the group if (1) the Euclidean distance between centroids is less than the set distance threshold:

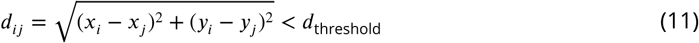

(2) the size ratio is less than the set size-ratio threshold, and (3) adding the new component would not exceed the maximum merged size. After groups are formed, components in each group are merged by pixelwise summation of their spatial images and the merged centroid is recomputed via a center-of-mass calculation:

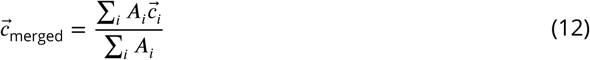

In the worst case, pairwise screening of *N* components is *O*(*N*^2^) with typically a few-hundred candidates; this pass is fast in practice.

### Step 4d: Initial Temporal Signal Extraction

Before any CNMF refinement, a conservative estimate of each candidate unit’s activity is needed to (i) verify that the segment actually carries calcium signal and (ii) provide stable seeds for subsequent updates. Thus, for every candidate region, a simple region-average trace by taking is formed, at each frame, the mean fluorescence of all pixels inside that region’s mask:

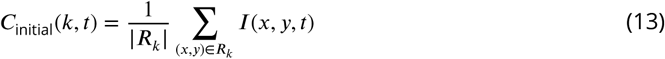

Stacking all masks as columns gives a sparse “mask matrix” that, when multiplied by the movie, produces all traces in one pass; each trace is then divided by its region area to yield comparable units. Additionally, the cost scales with the number of regions × pixels × frames, but because masks are sparse pixels are only iterated that belong to at least one region. The effective cost is proportional to (total masked pixels) × (number of frames), and thus, in a typical field (≈ 200 regions, ≈ 50 pixels each), that’s ∼ 10,000 pixel-sums per frame and is far smaller than processing the full image. Therefore, the data is streamed in time chunks (default 10k frames) to bound memory, with optional parallelism across regions/frames for seed initialization.

### Step 4e: Spatial and Temporal Matrix Initialization

With step 4e, the segmented footprints and their region-average traces are turned into consistent spatial/temporal matrices that downstream CNMF can use, while enforcing basic quality checks and reproducibility. Users can set (i) spatial normalization mode (none, max, L1, or L2), (ii) minimum component size (default 10 pixels), (iii) maximum number of components (0 = no limit), (iv) whether to skip any background component (ID 0), and (v) NaN/Inf checks. The surviving components are defined as the units and stack their 2-D footprints into a spatial tensor with shape [units, height, width] and their 1-D traces into a temporal matrix with shape [frames, units]. Coordinates (height/width) are inherited from the Step 3b template to keep all downstream arrays aligned. If spatial normalization is enabled, each unit’s footprint is rescaled by the chosen norm (max, L1, or L2):

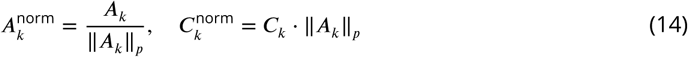

and its corresponding temporal trace is inversely rescaled so that the implied reconstruction magnitude (spatial × temporal) is preserved. Lastly, the cost is linear in terms of number of units and their pixel extents (stacking and per-unit normalization), plus lightweight I/O and logging. With a few hundred units, this step is quite fast in practice.

### Step 4g: Temporal Correlation Merging

For step 4g, MPS performs a single-pass, correlation-guided merge of candidate components using the cleaned spatial/temporal matrices from Step 4f. The GUI exposes four controls: (i) temporal correlation threshold (default 0.75), (ii) spatial overlap threshold (fractional, default 0.30), (iii) maximum merged size in pixels (default 5000), and (iv) an optional cap on the number of components to consider (0 = no cap). The temporal matrix is materialized to a dense array (*K* ×*T*) and computes a full Pearson correlation matrix with np.corrcoef:

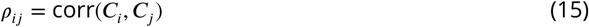

The overlap is computed using the overlap (Szymkiewicz–Simpson) coefficient (***Simpson*** (***1960***)): the count of intersecting pixels between two masks divided by the size of the smaller mask:

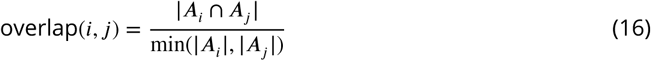

This metric equals 1 when the smaller mask is entirely contained in the larger and is generally ≥ IoU (Intersection over Union). Unlike IoU, which penalizes size asymmetry, the overlap coefficient is designed to capture inclusion and containment, making it especially well-suited for detecting redundant or duplicate cellular masks where one footprint may be fully nested within another. For surviving pairs, the algorithm grows a merge group by unioning any other components whose temporal correlation with either seed exceeds the threshold, provided (a) each proposed member overlaps spatially (by the same rule) with all current members and (b) the estimated union size of the group’s masks does not exceed the maximum-size limit. Components accepted into a group are marked processed to avoid reuse. All remaining components become singleton groups. Building the correlation matrix is *O*(*K*^2^ × *T*) in the worst case (after a full materialization of *C*); group formation uses set/overlap checks that are fast for “few-hundred” components. Default thresholds (temporal *r* = 0.75, spatial = 0.30) are chosen to be conservative and users can tighten/loosen them in the panel. However, these steps are for increasing convergence of the CNMF by providing a better starting point, thus it is better to include more components, than less in this situation.

### Step 5: Pre-CNMF Noise Estimation and Validation

Prior to CNMF refinement, MPS generates a pixelwise noise map from residuals of the background model and validates inputs for integrity and size-based plausibility. These checks establish component-specific noise levels used in the CNMF updates and ensure robust inputs for subsequent updates.

### Step 5a: Noise Estimation

To obtain a spatially resolved estimate of measurement noise prior to CNMF refinement, a per-pixel noise map is computed from the residuals of the background model. The input movie is first cropped and motion-corrected (Step 3a). The background is represented by a single spatial component (from Step 3b) and a corresponding temporal trace. A background movie is formed as their outer product, subtract it from the data, and obtain residuals that capture what the background model cannot explain:

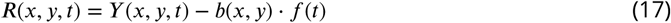

The per-pixel noise level is then estimated as the temporal standard deviation of the residuals, yielding a two-dimensional “noise standard deviation” image:

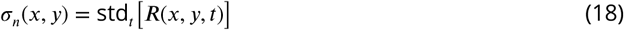

This map highlights which regions of the field of view fluctuate strongly after background subtraction, providing a more realistic measure of signal quality than a single global estimate. To account for heterogeneity in actively fluorescing regions, users are allowed to scale the noise inside “active background” areas by a multiplicative factor (default 1.5). Active regions are defined by thresholding the background component using either the global mean, global median, or a custom threshold. Inside the resulting mask, noise values are inflated accordingly. This adjustment helps prevent bright, slowly varying regions from being mistaken as low-noise simply because they dominate the background. For additional stability, the noise map can be Gaussian-smoothed, which reduces pixel-level spikiness while preserving broad spatial trends. Because the entire procedure is linear in both pixel count and frame count, and runs out-of-core with xarray/Dask, it scales efficiently to multi-hour recordings without exhausting memory.

### Step 5b: Validation and Setup

Before the alternating CNMF updates, an integrity check of inputs and (optionally) size-based component filtering is performed. Optional checks scan *Y*, *A*, *C*, and the noise map for NaN/Inf values and report global ranges; an estimated memory footprint (sum of nbytes) is logged to help users gauge resource demands. For footprint sanity checks, each unit’s area is computed as the count of strictly positive pixels in its spatial map; from these sizes summary statistics are reported and the counts falling below a user-selected minimum or above a maximum. If “Apply size filtering” is enabled, units outside the specified range are removed by indexing on the unit_id dimension.

### Step 6: Temporal Activity Update (CNMF Temporal Refinement)

Similar to CNMF implementations (***Lu et al.*** (***2018***), ***Giovannucci et al.*** (***2019***), ***Dong et al.*** (***2022***)), temporal activity is refined (*C*, plus sparse events *S* and background terms) while holding the current spatial maps (*A*) fixed. Importantly, proceeding with this temporal update first, unlike other CNMF implementations, drastically reduces total runtime since a cleaner *C* narrows the spatial search in Step 7. Additionally, all previous steps (initialization, seeding, merges, and noise estimation) exist as a warm start for the next CNMF updates to speed CNMF convergence and stabilize this temporal and next spatial pass.

### Step 6a–6b: Computation and Validation of Projected Fluorescence

Given the current spatial footprints and background model, Step 6a computes the residual activity tensor by fitting the movie as approximately the sum of *A* ⋅ *C* and *B*, then carrying forward the projections needed for deconvolution:

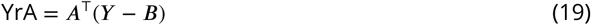

This leads to a marked increase in processing time by performing this calculation once instead of the current standard that performs the YrA computation for every component. In practice, the movie is reshaped into a two-dimensional array (frames × pixels), the background is optionally subtracted, values are cast to 32-bit floats to bound memory use, and any NaNs are replaced with zeros. The result is a dense projection of the movie onto the component footprints, which is then saved in chunked Zarr format for efficient downstream access. Additionally, a sanity-check was added to the YrA outputs before deconvolution by computing correlations and descriptive statistics and visualizing example traces and residual panels.

### Step 6c: Automated temporal parameter suggestion

While many existing tools require users to determine suitable parameters through trial and error, MPS provides automated suggestions to streamline setup, such as proposing conservative defaults for the AR model and sparsity based on Step 6a YrA, the Step 5a noise map, and a user-specified subset of components/frames. For reference, decay coefficients are estimated (AR order *p*, coefficients *g*) from autocovariance structure:

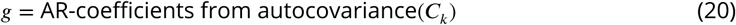

set a sparsity weight *λ* proportional to local noise, and (optionally) normalize traces. Selection can be random, best/worst SNR, or by size, and all suggestions are logged for reproducibility. These parameters are meant to be sensible starting points, not exact ground truth and the user is encouraged to tune them to their dataset.

### Step 6d: Update temporal components

In this step, each potential cell is assigned a calcium trace and corresponding spike train by solving a noise-aware sparse deconvolution problem that respects autoregressive calcium dynamics. For every unit, the method balances two terms: fidelity to the observed residual trace and sparsity of the inferred spikes:

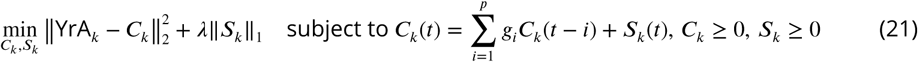

Non-negativity is enforced so that all outputs remain physically interpretable. The implementation uses CVXPY, an open-source Python library for formulating and solving convex optimization problems, with a formulation that explicitly introduces spike variables and ties them to calcium dynamics through an autoregressive operator, guaranteeing a convex and stable optimization (***Diamond and Boyd*** (***2016***)). Several design features make this implementation robust and scalable. Each trace is normalized, noise is estimated locally from the spatial footprint, and autoregressive coefficients are estimated adaptively in a way that ensures stability even on long or noisy recordings. Drift and offset terms are modeled explicitly, so slow baseline changes do not contaminate the spike train. If one solver backend fails, the system automatically falls back to alternatives, greatly improving reliability across datasets.

Instead of processing traces sequentially, the data is broken into overlapping temporal chunks that can be distributed across many workers. Within each chunk, all components are solved in parallel, and then chunks are reconciled to reassemble continuous outputs. This parallelization ensures that hundreds of calcium traces can be updated simultaneously, and thus, since the cost per trace is proportional to the number of frames, the end-to-end runtime scales linearly with the size of the dataset while achieving very high throughput in practice. Additionally, by performing the temporal update before refining the spatial footprints makes later spatial updates converge faster, reduces instability, and cuts down on redundant iterations.

Finally, the pipeline includes comprehensive convergence tracking. Metrics such as objective value, primal and dual residuals, solve time, and autoregressive coefficients are logged for each component. These can be saved, visualized, and inspected to verify that optimization is proceeding correctly. This level of transparency helps ensure that results are both reproducible and interpretable. Refer to step 8a for more information.

### Step 6e: Temporal quality control and filtering

Summary metrics are computed and remove obvious artifacts. Defaults include thresholds on total spike activity (min_spike_sum), variance of *c*_*i*_(min_c_var), and a minimal spatial-sum check to avoid degenerate *A*-*C* pairings. The UI provides quick inspection of individual components and saves both the full and filtered outputs with chunked, NumPy-backed export for fast reload in later steps. This temporal-first ordering materially decreases wall time for spatial refinement. With cleaner *C*, the localized regressions in Step 7 operate on smaller, better-conditioned tiles, require fewer passes, and parallelize across clusters without contention.

### Step 7: Spatial Footprint Update (CNMF Spatial Refinement)

Following the temporal update (Step 6), MPS updates spatial footprints while holding temporal activity fixed. Thus, after performing the temporal update first, cleaner temporal traces reduce ambiguity in how pixels and time are coupled - ensuring a faster spatial solve convergence, requiring fewer passes, and allowing for parallel execution across independent neighborhoods. The spatial update is designed for high parallel efficiency, so the workload is distributed across workers without heavy synchronization.

### Step 7a–7d: Dilation-Bounded Search, Neighbor Discovery, and Localized Regression

In Step 7a, each component’s support set is expanded conservatively by dilation with a small user-defined radius. This defines a growth mask Ω that bounds how far a footprint can spread, shrinking the search space and preventing spurious expansion:

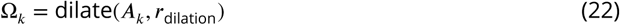

Step 7b builds a KD-tree (a k-dimensional, space-partitioning data structure for fast nearest-neighbor queries) over footprint centroids to identify spatial neighbors. Nearby or overlapping units are grouped into clusters, and for each cluster a bounding box Λ is computed. These clusters define disjoint tiles, allowing extremely parallel updates because clusters with no overlap can be processed independently.

Step 7c enforces conservative size bounds with a new per-component maximum growth parameter: each footprint can grow at most *γ* times its original support:

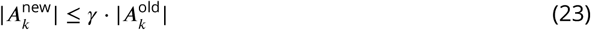

If an updated mask exceeds this cap, only the strongest pixels by coefficient weight are retained. This prevents runaway growth and guarantees that tile updates remain stable and independent.

Step 7d manages regression parameters and noise weights. Within each tile, a residual movie is formed by subtracting the influence of outside neighbors. A noise-weighted, nonnegative regression then updates footprints inside the dilation mask, combining L1 sparsity (to promote compact footprints) with L2 conditioning (to stabilize fits):

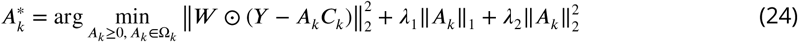

For single isolated units this reduces to a simple per-pixel non-negative least squares (NNLS) formula where CVXPY is the tool and NNLS becomes the problem structure.

### Step 7e–7f: Parallel Execution and Post-Update Merging

The spatial update is not a single “apply regression everywhere” operation; instead, it is a staged process designed to refine each footprint inside a bounded region, using sparse computation and explicit growth controls to keep results biologically meaningful and computationally efficient.

The function implements a multi-penalty sparse regression for every pixel inside a component’s dilation mask. Instead of fitting the entire field of view, the algorithm restricts attention to pixels flagged by the dilation mask Ω. This means that only a 5 to 20% fraction of the pixels in a tile are ever considered leading to an immediate several-fold speedup relative to dense updates. For each pixel in Ω, the local trace is extracted from the movie, normalized, and validated. Traces with extremely low variance or non-finite values are skipped to prevent numerical instability. Regressors are built from the temporal traces of the components in the cluster (and background signals). The solver then runs a multi-LASSO strategy: instead of choosing a single penalty, it evaluates a logarithmically spaced range of penalties, solves each with non-negativity enforced (using LassoLars), and collects statistics about sparsity, residuals, and support. These multiple solutions are then combined into a single coefficient vector, weighted by residual quality. This approach is robust to over or under penalization and provides a distribution of solutions. The pixel coefficient is accepted only if it clears an activity threshold. Coefficients below threshold are discarded, which ensures that noise fluctuations do not accumulate into spurious spatial spread. For accepted pixels, the algorithm logs residuals, sparsity levels, and support frequency across penalties, building a per-pixel statistical picture of confidence. These diagnostics are aggregated into maps (support frequency, residual map, coefficient distribution) that can be inspected for quality assurance.

A novel safeguard is the maximum growth factor. After solving all pixels in a component’s dilation mask, the number of active pixels is compared to the original footprint size. If the footprint has grown more than a user-defined multiple (default *γ* = 3×), the component is trimmed back. Specifically, pixels are sorted by coefficient magnitude, and only the top *γ* ⋅|*S*_old_| pixels are kept guaranteeing that components cannot explode in size due to overfitting or correlated noise. The growth cap is enforced consistently in two places: during per-component refinement and again when results are written back globally. The sparse design means that every step operates on a small fraction of the image, making the effective runtime proportional to the number of masked pixels rather than the entire tile. On typical datasets, this achieves an order-of-magnitude speedup relative to dense regression, while also yielding cleaner, more compact footprints. Together, these design choices transform the spatial update from a global, ill-posed problem into a sequence of small, well-posed, and bounded tasks. This is what makes it possible for MPS to refine thousands of footprints in multi-hour movies on a single workstation without exploding runtime or memory.

Step 7f finally handles post-update reconciliation. Overlaps are recomputed, and units with high similarity are merged. Statistics on area, compactness, and growth-cap enforcement are logged, and results are written back with full provenance. Because clusters were already defined to minimize contention, this merging stage is lightweight and can be parallelized as well.

Overall, ROI cropping in Step 3, watershed seeding in Step 4, noise-aware validation in Step 5, and temporal-first refinement in Step 6 all serve to make Step 7 a local, bounded, and highly parallel task. The dilation masks, cluster tiles, noise-weighted regression, and growth-cap innovation together transform one large ill-conditioned regression into many small, well-posed problems that can be solved side by side. This design reduces analysis time and memory pressure, improves demixing, and most importantly guarantees reproducibility across runs.

### Step 8: Final Temporal Refinement and Output

After spatial updates, MPS performs a final temporal pass using refined footprints to improve spike inference and calcium trace accuracy. Results are then exported for curation in the integrated Data Explorer, providing a consistent and reproducible endpoint for analysis.

### Step 8a–c: Re-projection, Temporal Updates, and Final Filtering

After spatial refinement (Step 7), the projection of the movie is recomputed onto the latest footprints to obtain an updated YrA (the per-component fluorescence obtained by multiplying *A*^⊤^ with the movie *Y*) exactly the same as Step 6a. In Step 8b, with YrA recomputed from the refined footprints, a last pass of temporal inference is performed. As in Step 6, each component’s trace *c*(*t*) is modeled with an auto-regressive (*p*) calcium dynamics prior and an L1 penalty on the innovation (spike) sequence, subject to nonnegativity. When a Dask client is available, Step 8b dispatches independent temporal chunks to workers to reconcile overlapped windows to avoid edge artifacts. The last sub-step assembles results for analysis and sharing. The UI loads any saved parameters (e.g., minimum footprint size, minimum SNR, minimum correlation, whether to include metadata, and the compression level), checks which spatial and temporal arrays are present, and detects whether spike estimates and background components are available from prior steps. On completion of the export, MPS celebrates with a small “fireworks” animation implemented in Tkinter. When the animation window is closed, MPS immediately offers a modal “Processing Complete” popup guiding the user to open the Data Explorer or continue elsewhere - so the celebration doubles as a clear hand-off into curation.

The final temporal pass repeats the same objective as the earlier update but with cleaner *A*, so both deconvolution and spike inference are better conditioned. In other words: *c*(*t*) is estimated that (i) explains YrA, (ii) is nonnegative, (iii) follows an AR(*p*) calcium evolution, and (iv) is sparse in its increments. Because footprints are now demixed and localized, YrA is less contaminated by neighbors, so the AR fit stabilizes and fewer iterations are needed. Moreover, chunked time windows with slight overlap allow linear-time passes over *T* while avoiding boundary artifacts when merging. The Step 8 implementation uses the same utilities (process_temporal_parallel, merge_temporal_chunks, defaults via get_default_parameters) as Step 6 to keep behavior consistent across passes.

### Interactive Data Explorer and Manual Curation

The Data Explorer is a post-processing application built explicitly for manual quality control by users. Its purpose is to let a trained human verify that each extracted footprint corresponds to an anatomically plausible ROI and that its calcium trace exhibits visible transients in the raw field - before results are accepted for downstream analyses. It is not an automated adjudicator; rather, it’s the standardized place where an expert inspects, accepts, or rejects components and documents those decisions.

The Explorer launches as a dark-themed app with a split layout: left is an interactive *A* view; right is a stack of per-neuron *C* traces. One can load A.npy and C.npy independently; the tool auto-detects and converts input shapes so that *A* is handled internally as (*H*, *W*, *U*) and *C* as (*U*, *T*). Clicking on a neuron in the *A* view both highlights its footprint and moves its trace panel to the top of the right stack, surfacing key units for rapid inspection. During playback, the *A* view is modulated frame-by-frame by the current activity level *C*[*t*], effectively “blinking” active cells so an expert can visually link transients in the trace to the underlying image. Default playback is 10 fps, adjustable from the menu. The right-hand trace pane shows each selected neuron in its own axis, with a movable time cursor and play/pause controls. Window length (seconds), playback step (seconds per tick), number of traces shown, and FPS are all configurable so an expert can tune the inspection workflow to the dataset. In the *A* view, the app can overlay neuron IDs, display a static max projection for orientation, and, when a neuron is clicked, report its estimated pixel area at a chosen fraction of the footprint’s peak intensity. These features are designed to answer the specific questions a human curator asks: “Does this footprint have a plausible size and shape?”, “Do the largest events in the trace manifest as visible fluorescence in the movie?”, and “Is this unit redundant with a neighbor?”

Curation actions are explicit and reversible. Cells can be deleted via the menu, and an Undo stack restores mistakenly removed units in place. The app maintains an “unsaved changes” state, and saving prompts the user to choose a destination and filenames; *A* and *C* are written as edited NumPy arrays (float32), leaving the original outputs untouched. Session metadata (e.g., animal & session labels, cache/export paths) is displayed at the top so curated files are clearly tied to the analysis run. These guardrails ensure that expert decisions are captured deliberately and reproducibly. To keep inspection fluid on long recordings, the trace view thins the x-axis to ∼ 1-Hz visual density by default, and playback uses a lightweight exponential moving average for smoother activity-weighted *A* rendering. The *A* composite is normalized for display and can be blurred slightly for legibility without altering the underlying data. Together, these choices favor fast human judgment without sacrificing fidelity.

After Step 8 produces final *A* and *C*, a user opens them in the Explorer, plays through the movie while watching selected traces, and prunes dubious units (vascular structures, edge artifacts, low-SNR remnants, duplicates). The curated *A*/*C* pair is then saved and carried forward to registration or behavior alignment. By centralizing this last mile of manual QC into a purpose-built tool, rather than ad-hoc scripts, the pipeline ensures that expert judgment is consistently applied and documented across datasets.

### Parallel Computing Implementation and Scalability Considerations

A cornerstone of MPS is the ability to handle extremely large movies with millions of pixels per frame (*M*) and 100,000 frames or more (*T*) on a single workstation without requiring a cluster. Every processing-expensive step is written to run on chunked arrays with out-of-core execution through Dask and vectorized NumPy/BLAS kernels, so memory use is bounded by the chunk size rather than the full dataset.

Time-chunked execution ensures that preprocessing and CNMF (Steps 2, 6, and 8) operate in windows of 1,000–5,000 frames at a time. Each window is read, processed, and cleared before the next begins. As a result, increasing *T* increases wall-clock time rather than peak RAM. On-demand spatial tiling is used only when necessary. If the field of view or number of units makes a single spatial pass impractical, MPS splits the update into non-overlapping tiles by dilating footprints and clustering nearby units. Only footprints that cross borders receive small overlaps, avoiding the reconciliation overhead of always-overlapping patches.

The scheduler relies on Dask’s distributed engine with a small pool of local workers (default 8). Large intermediates such as partial *A*^⊤^*Y* blocks are memory-mapped or written to disk, and tasks are short-lived to prevent buffer buildup avoiding the “giant graph” stalls that often occur in long notebooks. For example: 1) Motion correction uses FFT-based correlation in windows. 2) Temporal updates (Steps 6 and 8) run independently across units and windows. 3) Spatial updates (Step 7) are restricted to dilation masks within each tile and solved with noise-weighted non-negative regressions. Since the number of components *K* is much smaller than *M* × *T*, runtime is still dominated by the data size.

Theoretical complexity can be summarized as *T* × *M* log(*M*) for motion correction, *M* × *T* for per-frame preprocessing, *a* × *K* × *T* for trace extraction, *K*^2^ × *T* for correlations and merges, and (*M* + *T*) × *K*^2^ for NNDSVD. In a pessimistic bound with full frames, the motion term dominates and predicts ∼9–10h for a ∼3h video. In practice, Step 3 cropping reduces the effective pixel count *M* by 55–70%. This alone yields a 2.2–3.3× speedup across nearly every step. Combined with moderate *K* values (hundreds rather than the upper bound of 800) and the near-linear scaling of parallel workers, the actual wall time falls in line with about ∼1.3–2.6h for ∼ 95,000-frame runs. As an example, consider a 500 × 500 field of view, about 2.5 × 10^5^ pixels. With 100,000 frames and 800 components, the pessimistic calculation gives roughly 4.5 × 10^11^ operations from motion correction and another 2.2 × 10^11^ from NNDSVD, suggesting ∼ 9h. After a 60% crop the pixel count drops to 1.0 × 10^5^, which cuts those terms by a factor of ∼ 2.5, immediately bringing runtime to ∼ 3.5h. With parallel scaling across 8 workers and slightly smaller *K*, the observed ∼ 2h total for many datasets matches expectation.

Even though the workstation used for benchmarking had ∼ 512GB RAM, a hard cap of ∼ 200GB was enforced to emulate ∼ 256GB-class machines and keep usage predictable. Competing pipelines either required manual chunking and stitching or failed on multi-hour runs due to graph bloat or memory exhaustion, whereas MPS processed entire sessions in a single pass with bounded memory and continuous progress reporting.

### Statistical Analyses

Metrics were computed per recording and summarized across recordings with robust statistics. Baseline reduction (%) was calculated as one hundred times the difference between the pre- and post-values divided by the pre-value, where the measure is the median fluorescence over all pixels and frames. Cell-to-surround contrast was quantified as the fold-change in robust dispersion comparing post to pre. Rigid motion was estimated using frame-to-template cross-correlation; residual shift was defined as the median magnitude of per-frame displacement with interquartile range (IQR). Long-term drift, expressed in pixels per hour, was the slope from an ordinary least-squares fit of displacement versus time. Erroneous frames were flagged by heuristic detectors for line-split patterning, excessive blur, and saturation spikes. The effect of quality control was measured as the change in frame–template correlation, expressed in standard-deviation (SD) units relative to the pre-QC distribution.

For cropping and dimensionality reduction, metrics were computed on a per-session basis and summarized across sessions using robust statistics, including medians and interquartile ranges. Pixel load reduction was defined as the percentage decrease in active pixels per frame before and after cropping, while changes in runtime and memory usage were expressed as multiplicative speedup factors and percent reductions relative to the uncropped baseline. Variance explained was calculated as the cumulative fraction of total variance captured by truncated NNDSVD at a given rank *K*, and the elbow point was determined as the smallest *K* at which the marginal gain in variance explained dropped below one standard deviation of the pre-elbow slope. Iteration counts for CNMF initialization were compared between NNDSVD-seeded runs and both random and unconstrained least-squares baselines, with differences expressed as percent reduction in iteration number.

For cell detection, initialization, and pre-CNMF validation, component counts and areas were measured at each stage of the pipeline, beginning with watershed segmentation and progressing through mask merging, duplicate removal, and validation. Oversegmentation ratio was defined as the number of watershed masks divided by the number of curated components. Duplicate merging was quantified as the proportion of candidate pairs with both high temporal correlation (*ρ* > 0.8) and spatial overlap (IoU > 0.5) that were collapsed into a single component. Noise estimation was performed by computing pixelwise residual variance after subtracting a low-rank background, with optional Gaussian smoothing to stabilize variance estimates; for each component, noise level was summarized as the mean residual standard deviation across its pixels. Validation included checks for NaN and Inf values, size thresholds based on 1.5× the interquartile range of the component size distribution, and removal of invalid traces. The impact of quality control was quantified as the change in frame–template correlation after excluding erroneous frames, standardized in units of the pre-QC distribution. All analyses were conducted within individual sessions before aggregation across sessions to avoid distortion from outliers.

All statistics were calculated on a per-session basis and then summarized across sessions, with the session as the unit of analysis. Component counts before and after temporal quality control were compared using paired tests: normality was checked with the Shapiro-Wilk test and since the data were normally distributed, a paired *t*-test with within-subjects effect size was used. Percent rejection was reported as the average percentage of components removed across sessions. Signal-to-noise improvement was computed within each session as the mean ratio of denoised to raw traces, expressed in decibels, and compared using the same paired testing framework. Noise power was estimated from the median absolute deviation of the trace derivative, while signal power was taken as the variance of the trace. Decay times were derived from the autoregressive parameters and summarized with mean, median, and range.

Spatial refinement was evaluated per session and summarized across sessions. Component counts before and after refinement were compared with the same paired testing framework as above. Spatial organization was quantified with nearest-neighbor distances of component centroids, calculated using KD-trees, and summarized by the session-level mean and median. Density maps were generated to visualize local packing of components. Demixing quality was assessed by comparing temporal correlations among spatially overlapping neighbors before and after refinement. Morphology was summarized with measures such as compactness (perimeter squared divided by area) and centroid jitter across iterations; pre- and post-refinement values were compared per session with paired tests and effect sizes. When manual masks were available, specificity was quantified with intersection-over-union scores, reported as median and interquartile range per session.

## Acknowledgments

Funding for this work and for the authors’ research effort was provided by the National Institute on Drug Abuse (T32DA007278, R01DA052618, and R00DA052571) and by the Puget Sound VA Research and Development Pilot Grant.

## Notes

### Competing Interest Statement

The authors have declared no competing interest.

